# Unraveling cysteine deficiency-associated rapid weight loss

**DOI:** 10.1101/2024.07.30.605703

**Authors:** Alan Varghese, Ivan Gusarov, Begña Gamallo-Lana, Daria Dolgonos, Yatin Mankan, Ilya Shamovsky, Mydia Phan, Rebecca Jones, Maria Gomez-Jenkins, Eileen White, Rui Wang, Drew Jones, Thales Papagiannakopoulos, Michael E. Pacold, Adam C Mar, Dan R Littman, Evgeny Nudler

## Abstract

Forty percent of the US population and 1 in 6 individuals worldwide are obese, and the incidence of this disease is surging globally^1,2^. Various dietary interventions, including carbohydrate and fat restriction, and more recently amino acid restriction, have been explored to combat this epidemic^3–6^. We sought to investigate the impact of removing individual amino acids on the weight profiles of mice. Compared to essential amino acid restriction, induction of conditional cysteine restriction resulted in the most dramatic weight loss, amounting to 20% within 3 days and 30% within one week, which was readily reversed. This weight loss occurred despite the presence of substantial cysteine reserves stored in glutathione (GSH) across various tissues^7^. Further analysis demonstrated that the weight reduction primarily stemmed from an increase in the utilization of fat mass, while locomotion, circadian rhythm and histological appearance of multiple other tissues remained largely unaffected. Cysteine deficiency activated the integrated stress response (ISR) and NRF2-mediated oxidative stress response (OSR), which amplify each other, leading to the induction of GDF15 and FGF21, hormones associated with increased lipolysis, energy homeostasis and food aversion^8–10^. We additionally observed rapid tissue coenzyme A (CoA) depletion, resulting in energetically inefficient anaerobic glycolysis and TCA cycle, with sustained urinary excretion of pyruvate, orotate, citrate, α-ketoglutarate, nitrogen rich compounds and amino acids. In summary, our investigation highlights that cysteine restriction, by depleting GSH and CoA, exerts a maximal impact on weight loss, metabolism, and stress signaling compared to other amino acid restrictions. These findings may pave the way for innovative strategies for addressing a range of metabolic diseases and the growing obesity crisis.

## Main Text

The pioneering work of William C. Rose in 1937 revealed nine essential amino acids (EAAs): histidine, isoleucine, leucine, lysine, methionine, phenylalanine, threonine, tryptophan, and valine^11^. Notably, cysteine also behaves as an essential amino acid in animals with mutations in either cystathionine-γ lyase (CSE/CTH/CGL, here after referred to as CSE) or cystathionine-β synthase (CBS), enzymes of the trans-sulfuration pathway (Fig. 1a)^12,13^.

**Fig. 1:**
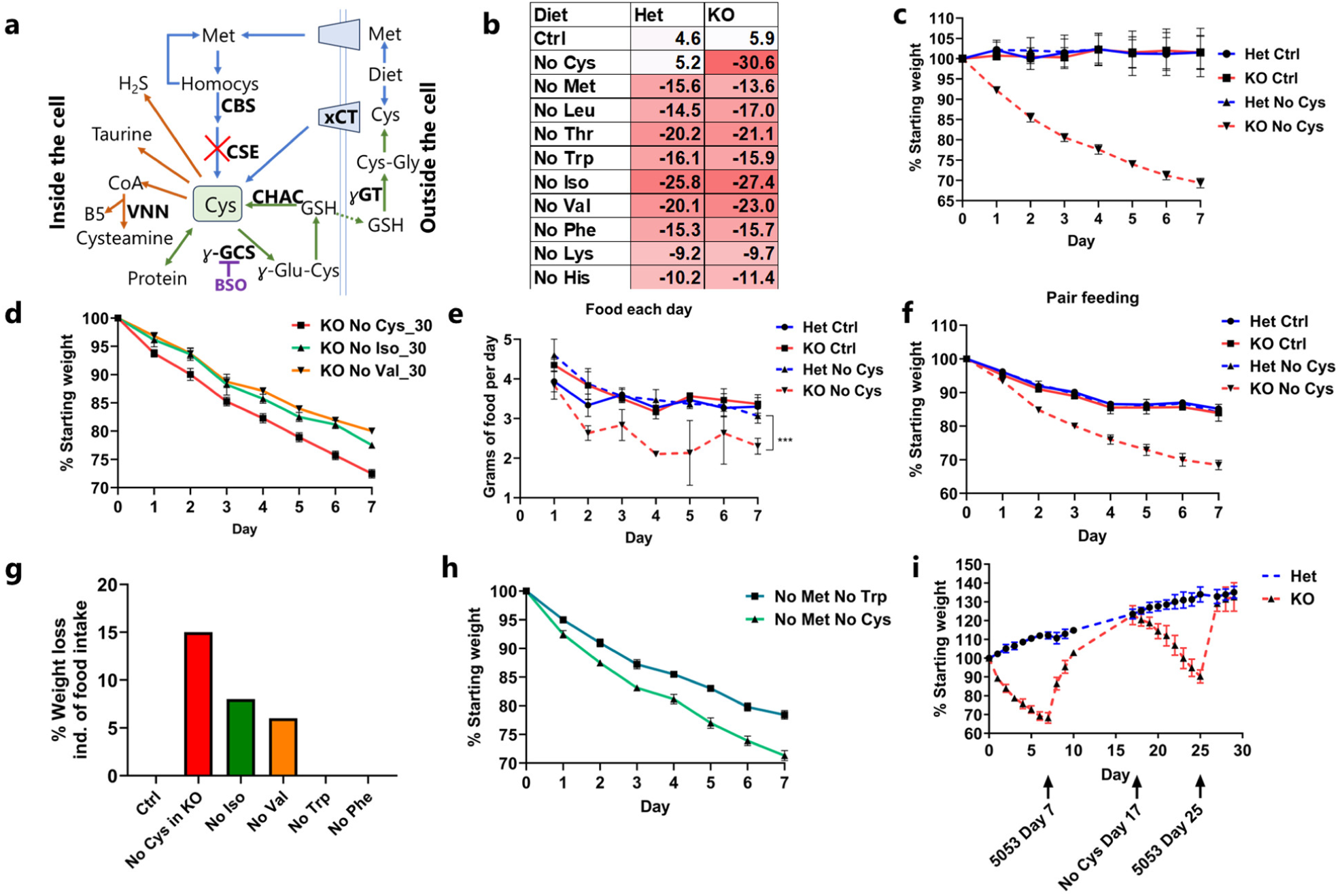
Cysteine deficiency drives rapid weight loss. **a**, A simplified cartoon demonstrating the pathways for Cys synthesis (blue) and consumption (reversible – green and irreversible – brown). CSE deletion marked by red cross. GSH synthesis inhibitor BSO shown in purple. **b**, Average percentage weight loss (from starting weight) with removal of each essential amino acid and cysteine in male Het and KO mice after 1 week (n>=4 for each group). **c**, Daily weight curves of male Het and KO mice fed control (Ctrl) or No Cys diets at 22 °C housing (n=4). **d**, *Cse* KO weight curves of mice deprived of isoleucine, valine, or cysteine at 30 °C (n=4) **e**, Daily food consumption of Het or KO mice with Ctrl or No Cys diets (n=3). **f**, Caloric restriction of 2.1 g/day (CR) of male Het and KO mice with control 5CC7 and No Cys diets (n=4). **g**, Average percentage weight loss unaccounted for by reduced food consumption (including data from PMID: 22016194 for Iso and Val, 34800493 for Trp and Phe)^14, 17^ **h**, Weight of male B6 mice on CR with No Met No Trp compared to No Met No Cys (n=4). **i**, Weight curves of male Het or KO mice over cycles of No Cys diet versus standard chow 5053 (n=4).

Over the course of several decades, extensive research has explored the effects of removing individual EAAs, shedding light on their roles in metabolism, energy expenditure, weight, and fat loss^14–18^. While weight loss in the absence of multiple EAAs is often attributed to reduced food intake, branched-chain amino acids (BCAAs), including isoleucine and valine, induce additional weight loss that may be explained by their role in thermogenesis^14^.

Amino acid deprivation triggers the integrated stress response (ISR) via GCN2, which detects uncharged tRNAs and phosphorylates translation initiation factor eIF2α^19^. Phospho-eIF2α suppresses general translation while promoting the translation of the key ISR transcription factor, ATF4, and the expression of its downstream targets, including the important signaling proteins fibroblast growth factor 21 (FGF21), an endocrine member of the FGF superfamily, and growth differentiation factor 15 (GDF15)^20–22^. The secretion of FGF21 results in increased energy expenditure and fatty acid oxidation^23,24^. Elevated GDF15 levels lower adipose stores and induce an aversive dietary response^7,8,20^. Oxidative stress can also trigger the ISR through endoplasmic reticulum (ER) stress, mediated by PERK^25^. This stress response stabilizes the transcription factor NRF2, and, in various cell lines and tumor models, NRF2 and ATF4 are reported to function cooperatively, reinforcing responses to amino acid limitation and oxidative stress^25–29^.

The combined low methionine and cysteine diet is notable because it increases lifespan and protects against metabolic diseases in rodents^30^. Although ISR activation and FGF21 upregulation are critical for some of those health benefits, it is still not clear whether the canonical GCN2-eIF2α-ATF4 axis is absolutely required^31^. Moreover, global FGF21 knockdown only suppresses thermogenesis but not lipolysis in adipose tissue, indicating that other factors are involved^23,32^. Finally, it is unclear if the benefits of methionine-cysteine dual restriction are driven by methionine or cysteine.

Cysteine is not only a proteinogenic amino acid but is also the limiting intermediary metabolite in glutathione (GSH) biosynthesis (Fig 1a)^12,13,33^. Cysteine additionally plays a critical, though underappreciated, role, alongside pantothenic acid (vitamin B5), in the synthesis of coenzyme A (CoA)^34^. In this pathway, pantothenate is typically regarded as the limiting factor, with its phosphorylation accepted as the rate limiting step^35,36^. CoA is also considered to be extremely stable, as mice on pantothenic acid deficient diets for as long as 2 months do not show significant loss of CoA^37^.

With a scarcity of research that comprehensively compares the mammalian response to the deficiency of each individual essential (and conditionally essential) amino acid within a single study, and the growing interest in diets low in sulfur amino acids and BCAA, we embarked on an investigation to examine the weight loss effect of removing from the mouse diet each essential amino acid individually. Our findings revealed that cysteine deficiency induces the greatest weight loss compared with all other EAAs, resulting in a surprisingly rapid reduction of approximately 30% of body weight within only seven days. Notably, this dramatic weight loss is fully reversible and accompanied by the browning of white adipose tissue mass. Our experiments have elucidated a coordinated mechanism underlying this phenomenon, consisting of rapid ISR and OSR activation, increased GDF15 and FGF21, and a drastic drop in CoA levels resulting in metabolic inefficiency, thus offering insights for potential intervention in metabolic disease and body weight control.

### Cysteine deprivation induces rapid weight loss

We initiated our study to evaluate the extent of weight loss induced when each of the nine EAAs and cysteine was individually removed from diet in both *Cse* knockout (KO or *Cse*^−/−^) and *Cse* heterozygous (Het or *Cse*^+/-^) C57BL/6 mice. While we observed significant weight loss in response to isoleucine and valine deficiencies, cysteine deprivation in *Cse*^−/−^ mice led to the most substantial weight loss (20% in 3 days and 30% in 7 days) compared to all other amino acids tested (Fig. 1b, c, Extended Data Fig. 1a-e)^14^. This effect was particularly remarkable considering that mice possess substantial reserves of cysteine stored as GSH^7,33^. Cysteine-free (No Cys) diet induced weight loss exclusively in *Cse*^−/−^ mice, with no observable difference between heterozygous and WT animals, indicating that depletion of newly absorbed and synthesized cysteine is necessary to induce marked weight loss (Extended Data Fig. 1f). We also observed no differences in percentage weight loss with changing starting weight (Extended Data Fig 1g-j). Female mice displayed slightly lower weight loss on Day 1, a difference which remained constant (Extended Data Fig 1k).

Weight loss was completely prevented by supplementation of cysteine through either N-acetylcysteine or GSH (which is broken down to cysteine in the gut) (Extended Data Fig. 2a). We also examined whether H_2_S, a degradation product of cysteine, played a role by giving daily injections of GYY4137 (a slow H_2_S releasing donor), but it did not prevent weight loss (Extended Data Fig. 2b)^38^. Microbiota alterations did not explain the weight loss, as antibiotic-treated mice still experienced the same degree of weight loss and co-housing *Cse*^−/−^ and *Cse*^+/-^ mice did not affect the weight of *Cse*^+/-^ mice (Extended Data Fig. 2c).

Considering the known roles of isoleucine and valine in thermogenesis in brown adipose tissue^39^, we measured the kinetics of weight loss in mice maintained in a thermoneutral environment (30°C). We observed that weight loss was still most pronounced for cysteine deficiency, with only a 2.7% change from the 22°C condition (Fig. 1d).

Because diets deficient in essential amino acids induce food aversion behavior, we closely monitored the daily food consumption of *Cse*^−/−^ and control heterozygous mice fed a cysteine-free diet compared to control diet. There was a 30% reduction in daily food consumption in *Cse*^−/−^ mice on the cysteine-free diet, decreasing from 3.5 g to 2.4 g/day (Fig. 1e). There was no difference between *Cse^+/-^* mice on control and cysteine-free diets (3.4 g in both). This food aversion and resultant caloric restriction (CR) could independently lead to rapid weight loss. CR of 2.1 g/day resulted in only 15-16% weight loss in the control conditions (15% in *Cse^+/-^* control, 16% in *Cse^+/-^* cysteine-free and *Cse^−/−^* control), whereas the *Cse*^−/−^ mice on the cysteine-free diet lost 31.5% of their weight within one week (Fig. 1f). Thus, weight loss of at least 15% could not be explained by reduced food intake. In contrast, for isoleucine and valine, the amount of weight loss unexplained by reduced food intake was 8% and 6%, respectively, as reported previously^14^. For other EAAs, such as tryptophan and phenylalanine, the entire weight loss was accounted for by reduced food intake (Fig. 1g)^18^, further emphasizing the unique effect of cysteine deprivation.

To determine if the weight loss was due to the accumulation of trans-sulfuration pathway intermediates such as homocysteine or cystathionine^12^, we provided *Cse*^−/−^ mice with a diet lacking both methionine and cysteine and compared the effect to that of the cysteine-free diet. The weight loss patterns were identical for the two diets, suggesting that neither homocysteine nor cystathionine were contributing to the weight loss (Extended Data Fig. 2d). Moreover, WT mice placed on a CR diet (2.1 g/day) devoid of methionine and cysteine lost approximately 30% of their body weight within a week as compared to those on a diet devoid of methionine and tryptophan that lost only 20% (Fig. 1h). This strongly suggests that benefits of sulfur amino acid restriction are primarily driven by cysteine rather than methionine limitation.

To investigate if the weight loss was due to poor nutrient absorption, at Day 3 of cysteine deprivation we compared stool metabolites from antibiotic-treated *Cse^+/-^* and *Cse^−/−^* mice. Antibiotic pre-treatment was conducted to minimize the potential impact of microbiota on differences in stool metabolites. We found no differences in amino acids, vitamins, glucose, or palmitic acid in the stool between *Cse^+/-^* and *Cse*^−/−^ mice, indicating no defects in nutrient absorption (Extended Data Fig. 3, Supplementary Table 1).

In light of the extremely rapid weight loss observed upon cysteine deprivation, we investigated whether reverting to a standard chow diet would result in weight gain. Strikingly, the mice regained about two/thirds of the weight loss within 2 days and fully recovered within 4 days (Fig. 1i). When the mice were switched back to a cysteine-free diet, they promptly resumed losing weight at a similar rate, which was again reversed immediately upon return to standard chow, emphasizing the high reversibility of cysteine deprivation-induced weight loss, which could be triggered multiple times without apparent detrimental effect.

### Cysteine deficiency induces selective fat burning and rapid browning

To further characterize the weight loss resulting from cysteine deprivation, we conducted metabolic and behavioral assessments with *Cse*^−/−^ and *Cse*^+/-^ mice in metabolic cages. Prior to placing mice in the cages, we subjected them to CR with a control diet (2.1 g/day) for 3 days to minimize the effects associated with CR only. After this period, we shifted the mice to metabolic cages and initially provided them the CR control diet for the first two days to establish the baseline profile and subsequently switched to the cysteine-free CR diet (Fig. 2a).

**Fig. 2:**
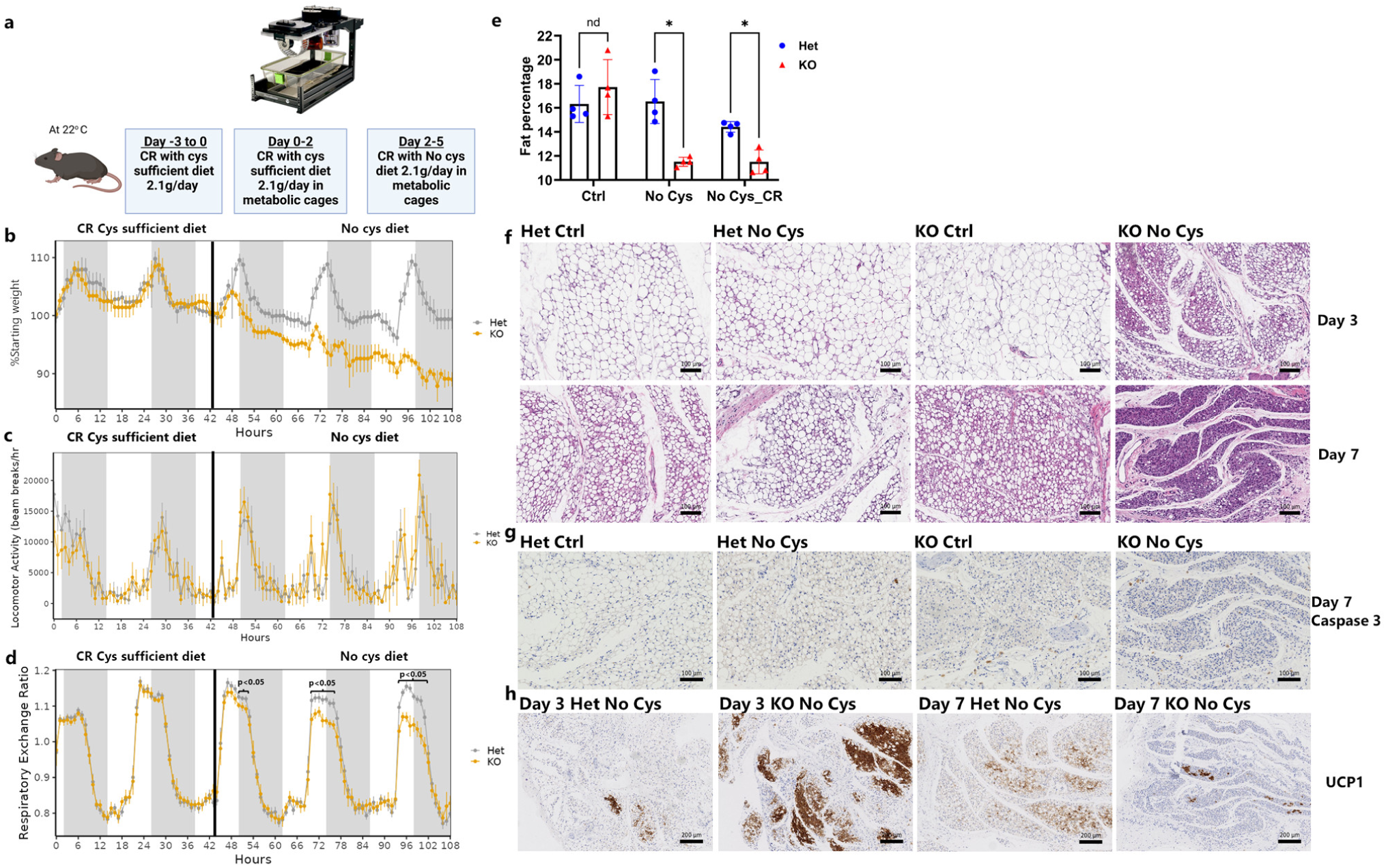
Cysteine deficiency drives rapid metabolic changes and loss of white adipose tissue mass. Metabolic cage profiles of male CR *Cse* Het and KO mice (n=4) for **a**, Experimental design **b**, Weight measurements **c**, Locomotor activity **d**, Respiratory exchange ratio **e**, DEXA comparing adipose tissue mass at ad-libitum and CR with cysteine-free (No Cys) diet and control (Ctrl) diets ad libitum at Day 7 (n=4). **f-h**, Representative images (n=4) of subcutaneous fat pads on CR. **f,** Male Het or KO on Ctrl or No Cys diets stained by H&E at day 3 and 7 imaged **g**, Immunohistochemistry of Caspase 3 on Day 7 and **h**, UCP1 on days 3 and 7 of cysteine restriction.

As anticipated, the shift to the cysteine-free diet in *Cse*^−/−^ mice immediately triggered weight loss, with a 10% decrease over 3 days, compared to 0% for *Cse*^+/-^ mice (Fig 2b).

Notably, there were no significant differences in locomotion and movement between the *Cse*^−/−^ and *Cse*^+/-^ groups under any condition, indicating that the weight loss is not attributable to increased physical activity in *Cse*^−/−^ mice, and that they do not display signs of lethargy (Fig. 2c).

The respiratory exchange ratio (RER), which shows whether mice are selectively burning fat or carbohydrate, was progressively reduced from Day 1 to Day 3 when *Cse*^−/−^ animals were shifted to a cysteine-free diet, suggesting increased usage of fat as fuel (Fig. 2d). To assess changes in total fat content, we performed DEXA scans on *Cse*^−/−^ and *Cse*^+/-^ mice on cysteine-free diet under both ad libitum and CR conditions. These scans revealed a substantial reduction in fat content in *Cse*^−/−^ mice on the Day 7 cysteine-free diet, even when the mice were on CR (Fig. 2e). No differences were observed at baseline between *Cse*^−/−^ and *Cse*^+/-^ mice when provided with the control diet.

To investigate whether this fat loss resulted from the loss of adipocytes or a uniform reduction of cellular lipid stores, we examined H&E-stained slides on subcutaneous white adipose tissue after CR on either cysteine-free or control diet. In *Cse*^−/−^ mice deprived of cysteine, there was significantly higher fat loss from individual adipocytes by Day 3, and by Day 7 there was near complete depletion of fat content throughout the tissue (Fig. 2f). Notably, all three control groups maintained significant fat content within each adipocyte on Day 3, which decreased slightly by Day 7. Caspase-3 staining on Day 7 revealed that despite significant fat loss there was no detectable cell death among adipocytes (Fig. 2g).

A closer examination of the H&E staining on Day 3 in *Cse*^−/−^ mice on a cysteine-free diet revealed a significant proportion of adipocytes that contained multiple small fat droplets instead of a single large droplet, resembling brown/beige adipose tissue. To investigate whether there was actual browning of the fat pad, we stained for UCP1. This revealed robust browning of white adipose tissue, on Day 3, at a pace much faster and more pronounced than previously reported with 4 weeks of CR (Fig. 2h)^40^. Additionally, there was rapid loss of visceral fat, which occurred even more quickly than the loss of large fat pads in *Cse^−/−^* mice on a cysteine-free diet. This was evident from the near-complete disappearance of fat surrounding mesenteric arteries by Day 3 (Extended Data Fig. 4a, b).

We did not observe significant differences in muscle (quadricep) histology, on Days 3 and 7, across any of the 4 groups (Extended Data Fig. 4c, d). Liver histology displayed no significant fat accumulation or apparent pathological changes (Extended Data Fig. 4e, f), in line with the reduced RER indicating fat utilization as an energy source in response to cysteine deficiency. The absence of appreciable differences may explain the lack of an effect on movement and the ease with which *Cse*^−/−^ mice on a cysteine-free diet can recover the lost weight.

### Transcriptional responses to cysteine deprivation

To gain insight into molecular responses triggered by cysteine deprivation, we conducted bulk RNA-sequencing of three key tissues: the liver (which exhibits the highest expression of *Cse*), muscle (the most abundant tissue), and adipose tissue (the tissue most impacted by cysteine loss). Recognizing that the lack of cysteine in *Cse*^−/−^ mice might trigger the ISR, we included a group exposed to a tryptophan-deficient diet to distinguish responses specific to cysteine deprivation and not general amino acid scarcity (Fig. 3a). To account for the potential gene expression changes driven by CR resulting from a cysteine-free diet, we employed a 3-day CR control diet and then shifted to either CR control, CR cysteine-free, or CR tryptophan-free (No Trp) diets for two days in both *Cse*^−/−^ and *Cse*^+/-^ mice.

**Fig. 3:**
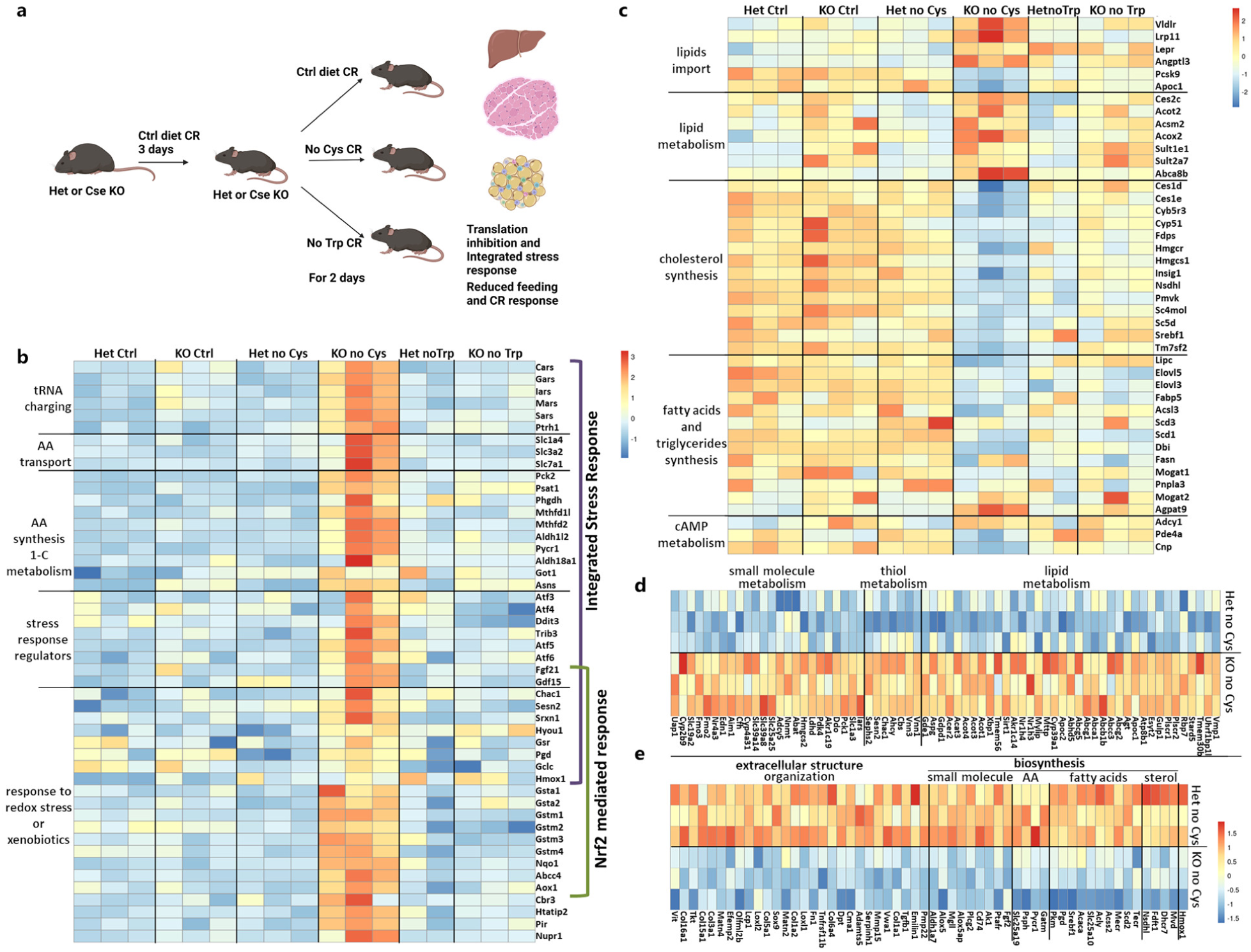
Changes in gene expression during cysteine deficiency compared to tryptophan deficiency. **a**, Experimental design for bulk RNA sequencing **b-c**, Liver bulk RNA-seq data represented as heatmap for genes related to ISR and OSR that are specifically upregulated in KO No Cys (AA, amino acids) (b), cholesterol and lipid synthesis and degradation (c) **d-e,** Heat map of genes that are specifically up- or down-regulated in epididymal adipose tissue in KO No Cys mice, including genes related to, lipid, thiol, and small molecule metabolism (d), and extracellular organization, biosynthetic pathways in small molecules, AA, fatty acids and sterols (e).

In the liver, there was distinctive transcriptional response specific for *Cse*^−/−^ mice on a cysteine-free diet (Supplementary Table 2). Gene ontology (GO) enrichment analysis of upregulated genes revealed prominent categories such as “cellular response to xenobiotic stimulus”, “small molecule” and “GSH” metabolic processes (Extended Data Fig. 5a). Closer examination of the “small molecule metabolic process” category revealed a strong upregulation of genes associated with ISR, including amino acid synthesis and one carbon metabolism (*Mthfd2, Pycr1, Asns, Psat1*), tRNA charging (*Sars, Cars, Gars, Mars*), amino acid transporters (*Slc7a1, Slc1a4, Slc3a2*), and various stress response genes (*Fgf21, Gdf15, Ddit3, Trib3, Atf5, Atf6*) (Fig. 3b, Extended Data Fig. 5b, c).

Although some genes typically associated with ISR (*Asns*, *Psat1*, *Pycr1* and multiple tRNA synthetases) were also upregulated in *Cse*^−/−^ mice on a tryptophan-free diet compared to *Cse*^+/-^ and *Cse*^−/−^ mice on control diet, their induction on cysteine-free diet was significantly more pronounced. This suggests that the magnitude of ISR activation depends on the specific amino acid deficiency (Fig. 3b).

Genes in the “cellular response to xenobiotic stimulus” and “glutathione metabolic processes” categories (*Nqo1, Gstm1-Gstm4, Gsta1, Gsta2, Srxn1*) are characterized by OSR regulated by NRF2. The concurrent activation of ISR (ATF4 signature) and OSR (NRF2 signature) in the liver suggests potential synergy (Extended Data Fig. 5a, Fig 3b), a phenomenon previously observed in cell lines and tumor models^26,27,29^.

The liver plays a central role in the metabolism of fatty acids (FA) and triglycerides, shifting between synthesis to breakdown in response to the fed and fasted states. As expected, several GO categories related to “cholesterol” and “lipid” metabolism were enriched among differentially expressed genes (Extended Data Fig. 5a, Fig. 3c). Genes associated with the import of lipid particles into the liver (*Vldlr, Lrp1l*) and key inhibitors of this process (*Pcsk9* and *Apoc1*) were up- and downregulated, respectively, in *Cse*^−/−^ mice on cysteine-free diet. This shift in gene expression suggests that the liver increases the import of very low-density lipoproteins (VLDLs) and low-density lipoproteins (LDLs), while reducing endogenous lipogenesis^41–43^. Furthermore, genes for sterol regulatory element-binding proteins (SREBPs), the master regulators of *de novo* lipogenesis (*Srebf1*) and cholesterol biosynthesis (*Srebf2*)^44^, and other genes associated with cholesterol biosynthesis, were significantly downregulated in *Cse*^−/−^ mice on the cysteine-free diet.

These changes are consistent with the suppression of *de novo* lipogenesis in the liver (Fig. 3c). Although some genes related to cholesterol and fatty acids biosynthesis were also downregulated in *Cse*^−/−^ mice on a tryptophan-free diet, the extent of downregulation was significantly less pronounced, underscoring a specific role of cysteine in the regulation of fat metabolism.

The transcriptional response to cysteine-free diet in muscle and liver tissue of *Cse*^−/−^ mice showed only partial overlap. In muscle, genes involved in the “response to oxidative stress” GO category were notably enriched. NRF2 (*Nfe2l2*) and its typical target genes (*Nqo1, Hmox1, Gclc, etc.*) were significantly upregulated in response to cysteine deficiency, indicating a robust induction of OSR in muscle (Extended Data Fig. 5d, e, Supplementary Table 3). The induction of several genes related to the import and catabolism of branched-chain amino acids (BCAA) (*Slc7a2, Idh2, Ivd, Bcat2*) suggests that muscle utilizes less glucose as an energy source, instead relying on amino acids that are not being utilized for translation. Although genes related to “extracellular structure organization” and “animal organ development” showed significant downregulation (Extended Data Fig. 5f), suggesting some structural rearrangements, muscle histology did not reveal any obvious pathological changes (Extended Data Fig. 4c, d). Interestingly, ISR was not upregulated in muscle, suggesting that the abundance of proteins in the tissue may prevent amino acid levels from falling below the threshold required to induce ISR at Day 2 (Extended Data Fig. 5g).

In epididymal adipose tissue, canonical ISR or OSR signatures were not observed (Extended Data Fig. 5h, Supplementary Table 4). On the contrary, some genes typically associated with ISR (*Pycr1* and *Psph*) and OSR (*Hmox1*) were significantly downregulated. To compensate for cysteine deficiency, genes such as *Chac1*, *Cbs* and *Ahcy* were upregulated. Furthermore, a significant suppression of SREBP-1 (*Srebf1*) and its target genes, such as *Scd2, Acly, Acaca,* and *Pgd,* indicated a shutdown of *de novo* lipogenesis in adipose tissue (Fig. 3d and e). Many genes related to lipid metabolism exhibited increased expression, including thioesterases (*Acot1-4*), implying increased release of free FAs from adipocytes. Although adipose tissue is not the major site for ketone body biosynthesis, genes in this pathway (*Acat3* and *Hmgcs2*) were upregulated. Additionally, there was a mild increase in *Ucp1* expression (Extended Data Fig. 5i).

### Rapid weight loss as a function of ISR and OSR

Because OSR-associated transcriptional changes were observed in both liver and muscle of cysteine-deprived mice, we explored whether there were alterations in GSH levels in these tissues. GSH levels can be influenced by dietary cysteine, and within just two days of cysteine withdrawal, there was a remarkable drop in GSH in the liver and muscle, but not subcutaneous adipose tissue, of *Cse*^−/−^ mice (Fig. 4a, Extended Data Fig. 6a-c). Furthermore, the upregulation of *Chac1* in the liver also likely contributes to the decrease in GSH levels (Fig 3b). This reduction was accompanied by the nuclear localization of NRF2 and increased NRF2-regulated NQO1 protein by Day 3 in livers of cysteine-deprived *Cse*^−/−^, but not *Cse*^+/-^ mice (Fig 4b).

**Fig. 4:**
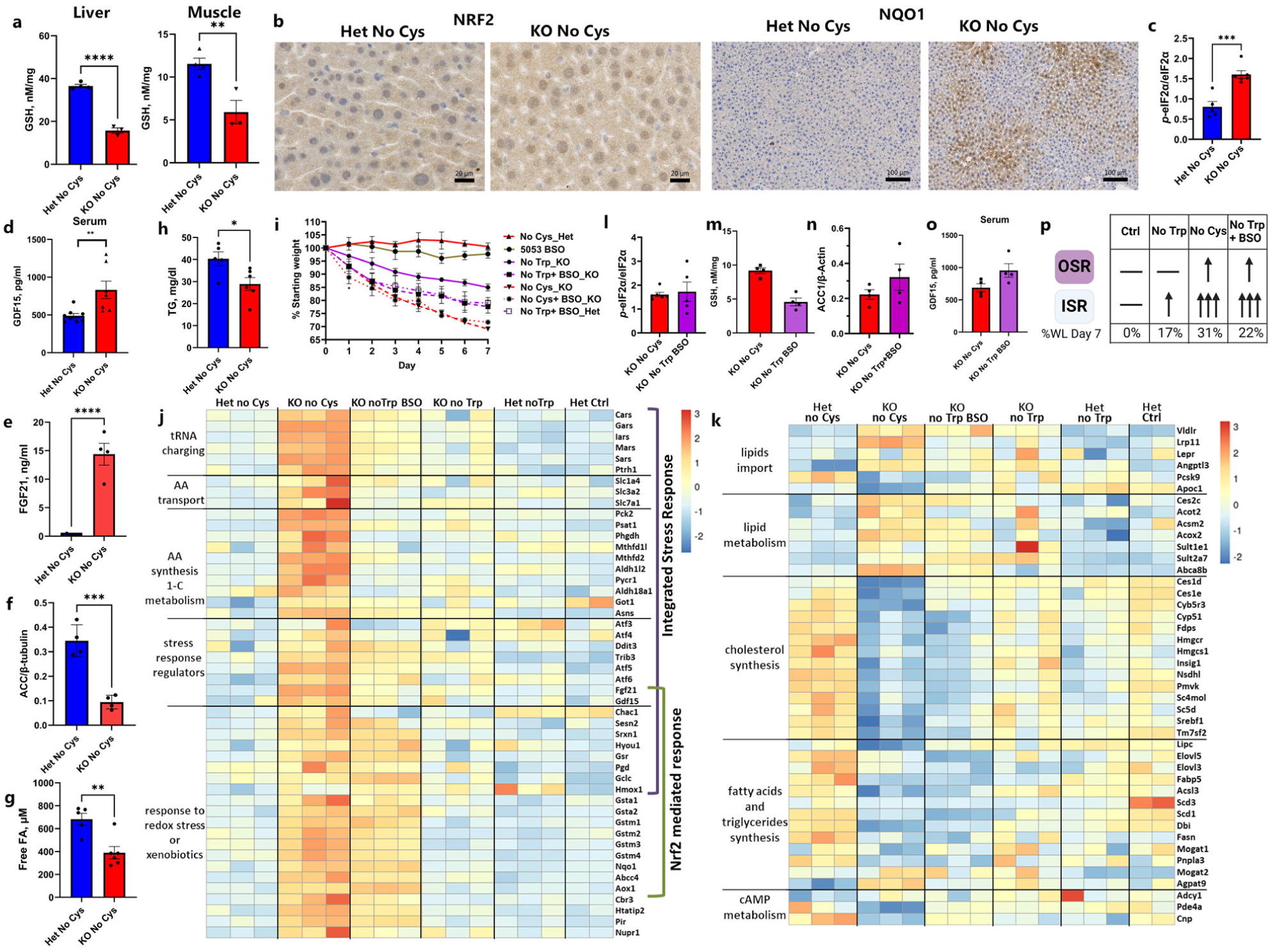
General EAA deficiency coupled with a deficiency in GSH partially phenocopies cysteine deficiency. **a**, GSH levels in liver and muscle of CR male Het and KO at Day 2 on a No Cys diet (n=4, n≥3). **b**, Representative IHC staining (of n=4) of NRF2 and NQO1 in liver of Het and KO mice on CR No Cys diet at Day 3. **c**, P-eIF2α/ eIF2α ratio in Het and KO mice on CR No Cys diet at Day 2 (n≥5). **d**, GDF15 and **e**, FGF21 serum level at Day 2 of CR No Cys diet (n≥7 for GDF15, n≥4 for FGF21). **f**, Acetyl-CoA carboxylase 1 (ACC1) protein levels normalized to β-tubulin in Het and KO No Cys at Day 2 on CR No Cys diet (n≥4).**g,** FFA and **h,** TG and serum levels at Day 2 of CR No Cys diet (n≥5 for FFA and TG)**i**, Weight measurements of *Cse* Het and KO mice on different diets (n>=4 in each group). **j,k,** Bulk liver RNA-seq data for multiple groups represented as heatmap for genes in Integrated Stress response and oxidative stress responses (g), and those related to cholesterol and lipid metabolism (h)**. l-o,** P-eIF2α/ eIF2α ratio in liver (l), liver GSH (m), ACC1 protein levels normalized to β-actin (n), and serum GDF15 (o) in calorie-restricted KO No Cys compared to KO No Trp + BSO mice at Day 2 (n≥4 for all). **p**, Summary of effects in KO mice by No Trp, No Cys and No Trp+BSO diets.

In light of the pronounced ISR signature in the liver, we further examined the level of phosphorylated eIF2α (P-eIF2α) and, as anticipated, found it to be increased on Day 2 in the livers of cysteine-free *Cse*^−/−^ compared to heterozygous mice (Fig. 4c).

Uncharged tRNAs activate GCN2 to increase eIF2α phosphorylation and activate the ISR. To verify ISR contribution to weight loss and food aversion we attempted to create *Gcn2 Cse* double knockout mice. However, 80% of such mice displayed hind limb paralysis and died with 8 weeks, thus preventing us from proceeding further. This phenotype suggests that *Cse^−/−^* mice, even on standard chow, exhibit a basal ISR driven by *Gcn2*, as evidenced by the mild increase in ISR genes such as *Asns* in the liver (Supplementary Table 2). Moreover, this experiment suggests that the ISR is absolutely required for adaptation to Cys deficiency, including weight loss.

GDF15 and FGF21 are both stress hormones associated with the ISR, and GDF15 has been shown to be induced during oxidative stress both *in vivo* and in cell culture^45,46^. Consistent with the upregulation of *Gdf15* and *Fgf21* transcription in the liver, within two days we detected a distinct increase in GDF15 and FGF21 in the serum of cysteine-free *Cse*^−/−^ mice, but not in the serum of Het mice or in mice fed other diets (Fig 4d, e, Extended Data Fig. 5b, c 6d, e). The observed increase in GDF15 may contribute to the heightened lipolysis and food aversion in cysteine-free *Cse*^−/−^ mice^8,10,46^. The increase in FGF21 also likely explains the observed increase in browning of adipose tissue.

Given the changes in GDF15 and FGF21, we decided to test if these stress hormones were also elevated in B6 mice upon methionine and cysteine dual restriction. As expected, we saw an increase in both GDF15 and FGF21 in the methionine and cysteine free diets but not methionine-free alone or control diet on both Day 3 and Day 7 (Extended Data Fig 7). To test if this elevation in GDF15 or FGF21 played a role in the weight loss, we gave *Gdf15* KO or *Fgf21* KO mice a methionine and cysteine free diet. We observed a significant decrease in weight loss in *Gdf15* KO mice on Day 1, but the difference decreased on Day 7 (Extended Data Fig 7f, g). This could likely be due to compensation by GCN2-mediated food aversion. There was also a significant attenuation of weight loss in *Fgf21* KO mice on Day 1, which became more marked over time (Extended Data Fig 7h, i). This is likely due to the role of FGF21 in browning of adipose tissue.

*Trib3*, was also upregulated in liver of cysteine-deprived animals. Acetyl-CoA carboxylase 1 (ACC1), the rate-limiting enzyme of fatty acid biosynthesis, which is negatively regulated by Trib3^47^, was markedly reduced in *Cse*^−/−^ compared to *Cse*^+/-^ mice on a cysteine-free diet (Fig 3b, 4f), further suggesting a reduction in *de novo* lipogenesis in the liver. Accordingly, we observed significantly lower triglycerides (TG) and free fatty acids (FFA) on Day 2 in the serum of the *Cse*^−/−^ compared to *Cse*^+/-^ mice on a cysteine-free diet (Fig 4g, h). Thus, increased utilization of fats and decreased synthesis contributes to rapid fat loss.

Given the significant changes in the liver following Cys restriction, we decided to test if restoring *Cse* expression in a liver specific manner can rescue weight loss. We administered intravenously 2×10^11^ AAV8 particles encoding either control EGFP or CSE driven by liver specific promoter TBG (TBG-EGFP or TBG-CSE), and waited 2 weeks before putting the mice on a cysteine-free diet. We observed a complete rescue of the weight loss, GSH levels in liver and serum TG and FFA levels, indicating liver expression of *Cse* is sufficient to prevent the weight loss (Extended Data Fig 8).

To confirm that weight loss in the setting of cysteine deficiency can be attributed to the induction of ISR and OSR, we fed mice a tryptophan-free diet and administered L-buthionine sulfoximine (BSO) at 25 mM in drinking water. BSO, a specific inhibitor of γ-glutamyl cysteine synthase, decreases GSH but not cysteine (Fig 1a). By Day 3, cysteine-free and tryptophan-free+BSO diets induced approximately 20 vs 18% weight loss in *Cse*^−/−^ mice, but by Day 7 the tryptophan-free+BSO group lost only an additional 4%, reaching 22% weight loss, while the cysteine-free group lost an additional 11%, resulting in 31% weight loss. This left an unexplained 9% difference in weight loss (Fig. 4i).

To investigate the potential cause of the remaining 9% weight loss, we compared the transcriptional response in livers of *Cse*^−/−^ mice on cysteine-free versus tryptophan-free+BSO diets. The transcriptional upregulation of ISR was slightly weaker in the tryptophan-free+BSO group compared to the cysteine-free group, notwithstanding the similar level of P-eIF2α (Fig. 4j and l). Genes associated with tRNA charging, amino acid transport, and key regulatory genes like *Fgf21* were only slightly increased in the tryptophan-free + BSO group compared to the tryptophan-free diet. However, genes associated with amino acid synthesis and one carbon metabolism remained similar to the tryptophan-free diet (Fig. 4j).

We found a remarkably similar upregulation of OSR in *Cse*^−/−^ mice fed either tryptophan-free+BSO or cysteine-free diets (Fig. 4k). This finding is logical considering the comparably low levels of GSH in the liver of *Cse*^−/−^ mice on tryptophan-free+BSO and cysteine-free diet (Fig. 4m). Moreover, inclusion of BSO significantly downregulated genes involved in cholesterol and fatty acids biosynthesis, as suggested in other studies involving GSH depletion^48^(Fig. 4k). On Day 2, ACC1 protein level was also downregulated by tryptophan-free+BSO (Fig. 4n) and GDF15 serum levels were similar in *Cse*^−/−^ mice fed either tryptophan-free+BSO or cysteine-free diets (Fig. 4o). These results indicate that in the liver, GSH content primarily controls the OSR and *de novo* fatty acid and cholesterol biosynthesis, while only certain aspects of the ISR are influenced.

Notably, CHAC1, the protein responsible for releasing cysteine from GSH, was not upregulated by BSO (Fig. 4k). This suggests that additional regulators may be activated, not by uncharged tRNA (ISR) or low GSH level (OSR), but rather by cysteine or its derivatives. Additionally, the presence of BSO in the cysteine-free diet did not further increase the weight loss in *Cse*^−/−^ mice (Fig. 4i) except on Day 1, likely due to faster GSH depletion.

### Cysteine depletion leads to rapid coenzyme A loss and metabolic inefficiency

Notwithstanding similar ISR and OSR responses, *Cse*^−/−^ mice on cysteine-free diet lost 9% more weight by Day 7 compared to those on tryptophan-free + BSO diets (Fig 4p). This result implies a role for another cysteine-containing molecule that is depleted. We considered Coenzyme A (CoA), a critical component of fatty acid metabolism, TCA cycle, and many other metabolic processes, since it is synthesized through the condensation of pantothenic acid (vitamin B5), cysteine, and ATP (Fig. 5a)^49^. Previous studies suggested that CoA is extremely stable and its levels cannot be easily altered. However, it was not known what would happen to CoA if cysteine was restricted. We observed a striking 30% reduction in total CoA levels in the liver and muscle by Day 2 and a 75% reduction in liver by Day 6 in *Cse*^−/−^ mice compared to *Cse*^+/-^ mice on a Cys-free diet (Fig. 5b-d).

**Fig. 5:**
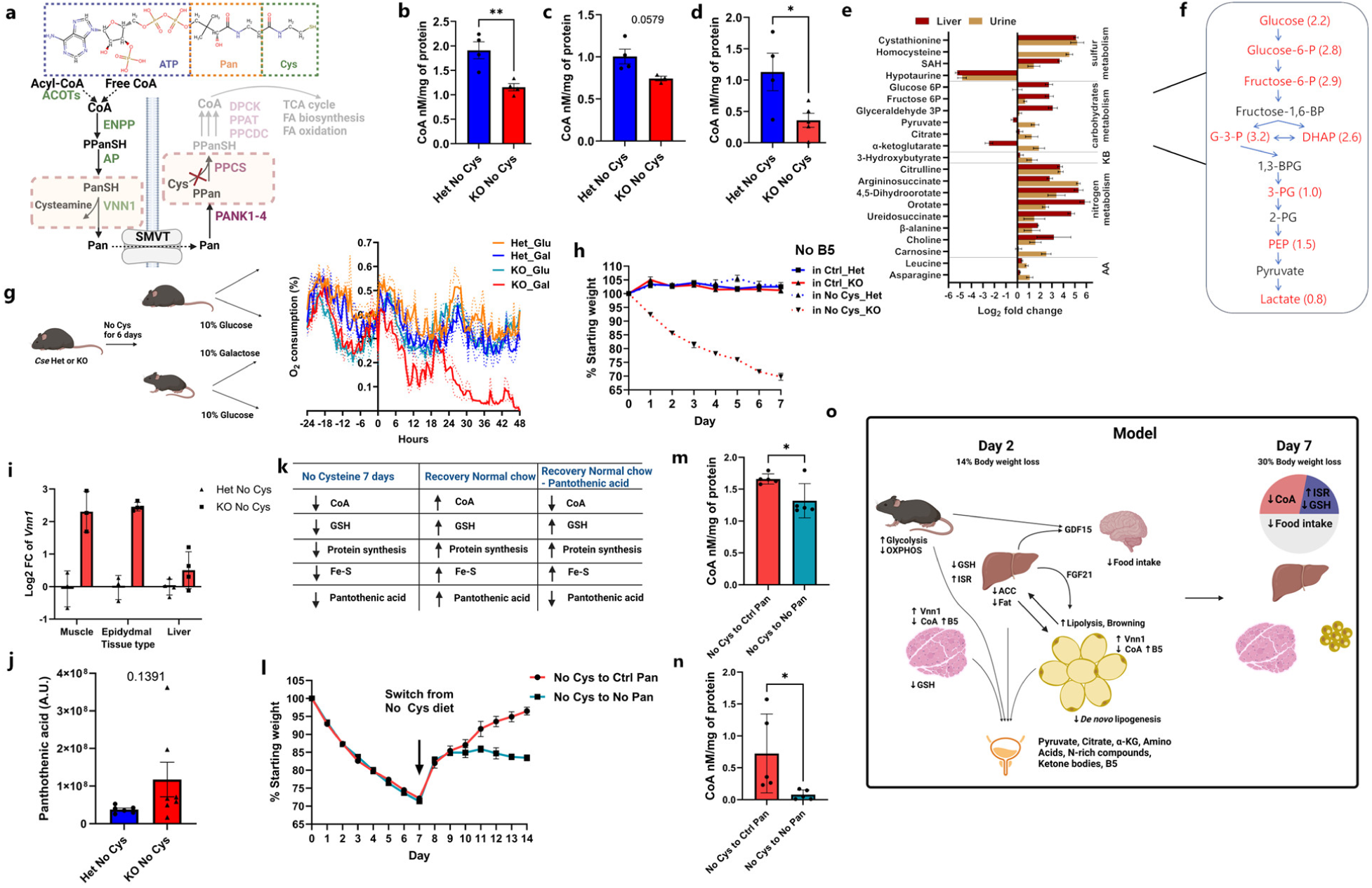
Cysteine deficiency leads to metabolic inefficiency by depleting CoA. **a**, Structure of CoA and pathways of CoA production and degradation. The red cross depicts how Cys deficiency would inhibit CoA biosynthesis and subsequent pathways involving it. **b-d,** CoA levels in liver (b) and muscle (c) at day 2 and liver at Day 6 (d) from *Cse* Het and KO mice on a No Cys diet (n≥3 for all groups) **e**, Select metabolite levels from urine and liver represented as log2 fold change between KO No Cys and Het No Cys (n≥6 for urine, n=3 for liver), AA: amino acids; KB: ketone bodies. Metabolites have *p*-values of less than 0.05 in either urine or liver except for 3-Hydroxybutyrate (*p*-value=0.0891). **f**, Liver glycolysis intermediate accumulation in KO compared to Het mice at Day 2 on CR No Cys. Red indicates up in KO with *p*-value<0.05. Number in bracket denotes log2-fold change (n=3). **g**, O_2_ consumption by mice provided Galactose or Glucose as sole carbon source after feeding No Cys diet (n=4). The experimental strategy is shown on the left. **h**, Weight measurements of male *Cse* Het and KO mice on No Pantothenic acid (No B5) diet or No Pantothenic and No Cys diets (n=4). **i,** *Vnn1* mRNA expression levels in muscle and epidydimal tissue from bulk RNA-seq data (as shown in Fig. 3a). **j**, Urine pantothenic acid levels in *Cse* Het and KO mice on CR No Cys diet at Day 2. **k,** Summary of expected levels of various cysteine-containing metabolites under different conditions **l**, Weight measurements of male *Cse* KO mice after 7 days on a No Cys diet followed by 7 days on either a control diet or a No B5 diet (n=5) **m**, Liver and **n**, Muscle CoA levels in mice on Day 14 of the experiment shown in **l**. **o,** Model summary of the effect of cysteine deficiency on metabolic pathways and weight loss.

Considering the central role of CoA in energy metabolism, its deficiency is expected to result in substantial abnormalities in cellular metabolism. We therefore examined the liver and urine metabolomes of *Cse*^−/−^ and *Cse*^+/-^ mice after 2 days on a cysteine-free diet. As expected, mass spectrometry analysis revealed an increase in cystathionine (upstream of CSE) and decrease in hypotaurine (downstream of CSE) in both liver and urine (Fig. 5e, Supplementary Table 5). There were marked increases in urine in the levels of pyruvate, citrate, and α-ketoglutarate (αKG), metabolites that precede steps involving CoA in the TCA cycle (Fig. 5e, Extended Data Fig. 9a, b). Furthermore, in the livers of *Cse*^−/−^ mice, there were notable increases in nearly all glycolysis intermediates (Fig. 5f, Extended Data Fig 9a).

CoA is also indispensable for the synthesis and degradation of fatty acids and amino acids. Our findings suggested that FA degradation and conversion to ketone bodies (KB) on the second day of a cysteine-free diet proceed unimpeded, as indicated by the observed increase in ketone bodies in the urine (Fig. 5e). Furthermore, the elevated levels of intermediates in the urea cycle indicate that dietary amino acids in the liver are degraded for glucose or energy production. This is supported by the elevated citrulline, arginosuccinate and intermediates of pyrimidine biosynthesis detected in urine, resulting in the wasting of intermediary metabolites (Fig. 5e, Extended Data Fig. 9a, c). Despite this, we saw elevated levels of multiple amino acids in urine, suggesting the diminished ability of amino acids to enter the TCA cycle or be used for gluconeogenesis due to low CoA (Fig. 5e, Extended Data Fig. 9).

To gain further insights into how carbohydrate metabolism changed with CoA depletion, we performed ^13^C-Glucose tracing in *Cse*^+/-^ and *Cse*^−/−^ mice. On Day 3 of a cysteine-free diet, we administered a bolus of U-^13^C-glucose (1.25mg/kg body weight) orally to 18h starved *Cse*^+/-^ and *Cse*^−/−^mice and studied the liver (after 45 minutes) and urine (after 2h) metabolomes (Extended Data Fig 10a, Supplementary Table 6). Consistent with inefficient conversion of pyruvate to Acetyl-CoA, there was a clear trend for elevated m+3 lactic acid in the liver of Cse-/- mice (Extended Data Fig 10b, c).

There was also a significant increase in urine of m+3 orotate, which is derived from pyruvate carboxylation to oxaloacetate, demonstrating another mechanism by which *Cse*^−/−^mice can waste carbon in the absence of CoA. (Extended Data Fig 10b and d). Additionally there was a significant increase in m+2 Creatine and its precursor guanidoacetic acid in both liver and urine of *Cse*^−/−^ mice (Extended Data Fig 10b and e-h). As *Cse*^−/−^ mice lost considerable fat, elevated creatine futile cycling could contribute to thermogenesis during the weight loss^50^. Curiously, we detected the upregulation of the cytoplasmic creatine kinase gene (*Ckm*) concomitant with the suppression of its mitochondrial isoform (*Ckmt*) in epididymal fat of *Cse*^−/−^ mice (Extended Data Fig 10i). We further observed the increased peak energy expenditure in *Cse*^−/−^ mice on cysteine-free diet along with higher total liver creatine level on Day 6 (Extended Data Fig 10j and k). Together these results are consistent with CoA deficiency compromising mitochondrial activity, necessitating cytoplasmic futile creatine cycling for thermogenesis.

Collectively these findings suggest that after 2 days of cysteine deprivation, the carbon skeletons from dietary glucose and amino acids are lost, appearing in the urine in the form of pyruvate, citrate, creatine, orotate, αKG, urea cycle intermediates, and pyrimidines, while lipid stores are utilized for ATP production. This preference for fat utilization, even when dietary glucose and amino acids are abundant, may contribute to the observed additional weight loss. Additionally, the increased excretion of acyl-carnitines in urine suggests futile attempts to recycle CoA (Extended Data Fig. 9d).

Consistent with systemic reduction of CoA, there was decreased basal O_2_ respiration in lymph node T cells from Day 7 cysteine-deprived mice, indicating compromised OXPHOS capacity (Extended Data Fig. 11a). Notably, this effect was specific to a cysteine requirement, as *Cse*^−/−^ mice receiving a tryptophan-free diet did not exhibit reduced basal O_2_ respiration (Extended Data Fig. 11b).

To further investigate the systemic requirement for cysteine-derived CoA, we restricted *Cse*^−/−^ mice to either 10% glucose or 10% galactose solution as the only energy source in their drinking water after 6 days on either a control, cysteine- or tryptophan-free diet (Fig. 5g). Unlike glucose, which can yield a net of 2 ATP through glycolysis, galactose provides no net ATP, and thus mice are unable to survive on galactose for an extended period if there is a general defect in TCA cycle entry. Within 24h of receiving only galactose as their carbon source, there was a marked decrease in O_2_ consumption only in the cysteine-free *Cse*^−/−^ mice. These mice required euthanasia by 42h, a situation that did not occur with control or tryptophan free-diets (Fig. 5g, Extended Data Fig. 11c, d).

As expected, no weight loss was observed with a B5-free diet, and there was no additional effect when B5 was removed from the cysteine-free diet. This underscores cysteine as the primary limiting factor for CoA biosynthesis (Fig. 5h). We then revisited our RNA-seq datasets to seek an explanation for the reduced CoA levels, as cysteine deficiency would only prevent new CoA production. One of the most significantly upregulated genes in both adipose and muscle tissues was *Vnn1*, a pantetheinase that degrades pantetheine into B5 and cysteamine (Fig. 5a)^49^. Although *Vnn1* is known to be triggered during starvation and fasting responses, enabling the diversion of CoA to the liver, we observed upregulation of *Vnn1* in cysteine-free *Cse*^−/−^ mice compared to *Cse*^+/-^ mice despite identical food intake (Fig. 5i)^49^. This increased *Vnn1* expression likely leads to enhanced degradation of CoA and the deficiency of cysteine prevents resynthesis of CoA, resulting in its insufficiency and subsequent loss of B5 in urine (Fig. 5j).

In mice deprived of cysteine or on a tryptophan-free diet + BSO, we also detected a slight but consistent decrease in expression of multiple genes associated with mitochondrial respiratory complexes. This may be an adaptive mechanism to mitigate ROS (Extended Data Fig. 12). The more pronounced mitochondrial defect in the absence of cysteine may be the consequence of the additional deficiency of CoA, resulting in greater metabolic inefficiency, and likely explains the additional 9% weight loss observed in *Cse*^−/−^ mice on a cysteine-free diet compared to those on a tryptophan-free diet + BSO.

There are multiple ways for Cys deficiency to induce metabolic inefficiencies. Beside depleting CoA, low Cys availability can compromise Fe-S clusters assembly, downregulate mitochondrial protein biosynthesis or disturb the redox balance through low GSH. To address the specific contribution of CoA, we relied on the fact that it must be degraded to pantothenic acid (vitamin B5) for transport between the tissues. This is suggested by the upregulation of *Vnn1* expression in peripheral tissues of *Cse^−/−^* mice on a cysteine-free diet (Fig. 5i). Moreover, some water-soluble vitamin B5 was lost in the urine (Fig. 5j). That implies that by Day 7 on a cysteine free diet, along with the depletion of other cysteine containing molecules, the *Cse^−/−^* mice would also have lower B5 levels across all tissues (Fig. 5k)

Indeed, when we reverted cysteine-deprived *Cse^−/−^* mice to a cysteine-sufficient but B5-deficient diet, they failed to gain as much weight as mice on the control B5 and cysteine-sufficient diet (Fig. 5l). Accordingly, GSH was fully rescued in the liver and partly in the muscle, while CoA levels were still significantly lower on the B5-deficient diet (Fig 5m, n, Extended Data Fig 13a, b). Furthermore, restoring B5 in drinking water after 7 days on the B5-deficient diet immediately rescued the difference in weight recovery (Extended Data Fig 13c). Taken together our results indicate that the loss of CoA contributes significantly to the metabolic inefficiency and weight loss on a cysteine free diet (Fig. 5k). The loss of CoA alone can lead to significant changes in metabolism, contributing to rapid weight loss and preventing weight gain, showing that it is a primary regulator of metabolic efficiency.

## Discussion

Our unexpected results indicate that cysteine deprivation triggers a global reprogramming of metabolic processes, culminating in rapid and readily reversible weight loss through the decline in adipose tissue lipid content, reduced lipogenesis and excretion of intermediary metabolites that cannot be effectively utilized. The results extend earlier studies that demonstrated an essential role of cysteine in *Cse^−/−^* mice and also reported weight loss, albeit at a much slower rate^12,13^. Additionally it was found that liver function, as determined by plasma levels of AST, ALT, and albumin, remained consistent between *Cse*^+/+^ and Cse^−/−^ mice even after 10 weeks on a 0.05% cysteine diet, as compared to the standard 0.4% cysteine diet^13^. Our results also show that cysteine rather than methionine mediates the benefits of methionine-cysteine dual restriction. Together, these findings suggest that targeted manipulation of CSE and dietary cysteine uptake could potentially offer a novel approach to induce rapid fat loss without adversely affecting skeletal muscle and other vital organs.

Due to its potent cytotoxicity^51^, cellular concentration of cysteine is the lowest compared to other amino acids^52^. In mammals most of dietary cysteine is absorbed by the liver and rapidly converted into much safer molecules - GSH and taurine^53^. When animals are exposed to a cysteine-free diet, intracellular cysteine is rapidly depleted due to protein, GSH, and CoA synthesis. Thus, a combination of low concentration and a high demand in multiple cellular processes likely explains an amplified ISR in response to cysteine compared to other amino acid deficiencies.

An intriguing aspect of our findings is the failure of millimolar levels of GSH in various tissues to prevent or even delay the weight loss by serving as a reservoir for cysteine within the cells. It appears that animals prefer to reduce metabolism by decreasing CoA levels and suppressing protein synthesis, rather than allowing rapid release of cytotoxic cysteine from GSH. The hypothesis finds support from studies with humans, in which elevated cysteine levels were closely associated with a range of pathological conditions, including obesity, neurological disorders, autoimmune and cardiovascular diseases^54–58^.

Accordingly, evidence from studies in rodents and nematodes has shown that restricting sulfur amino acids can extend lifespan^59–61^, further supporting the potential benefits of CSE inhibition. The decrease in GSH and CoA may be attributed to the futile attempts to salvage CoA between tissues. This is due to the increased degradation of muscle CoA into cysteamine and B5, which cannot be effectively reutilized in the liver due to the lack of cysteine. Cysteine breakdown into H_2_S could also play a role, although it is worth noting that supplementing with H_2_S did not rescue the weight loss. Consequently, this cascade leads to an increased loss of B5 in urine.

The upregulation of vanin (*Vnn1*) and rapid depletion of CoA levels throughout the body, leading to increased reliance on glycolysis for energy production, is another intriguing finding in our study. This decline in CoA levels occurs swiftly following cysteine deprivation, without any genetic defects in the direct pathway for CoA biosynthesis. The resulting CoA deficiency leads to pronounced lipolysis in adipose tissue due to a preference for utilizing fatty acids as the primary energy source, as evidenced by decreased RER. Previous studies have reported ketosis and the excretion of biomass in urine in diabetic models^62^. The utilization of fatty acids rather than carbohydrates when CoA levels are limiting could potentially be driven by lower Km for acyl-CoA synthetases compared to that for pyruvate dehydrogenase^63,64^. Thus, as CoA becomes limiting, lipid could become the preferred energy source leading to the lower RER observed. Our study demonstrates that cysteine deficiency-induced CoA loss results in metabolic reprogramming and inefficiency leading to the excretion of both glycolytic and TCA cycle intermediates, ketone bodies and other biosynthetic intermediates in the urine, contributing to the observed weight loss.

Both GSH depletion and ISR also suppress mitochondrial activity, reducing ROS production, and preventing the formation of new mitochondria^65,66^. Thus, the combined effect of ISR, GSH loss, and CoA deficiency contribute to overall mitochondrial inefficiency.

Our analysis of the liver RNA-seq data from cysteine-deficient mice revealed a robust ISR and OSR signature due to GSH depletion, a synergy previously reported in tumor cell lines^25,26,28^. Simultaneous activation of both pathways leads to rapid induction of GDF15, which was detectable within just two days. The role of GDF15 in inducing food aversion behavior likely explains the more pronounced reduction in food intake observed on Day 2 and Day 3 following the switch to a cysteine-free diet, as GDF15 requires time to accumulate^8,10,46^.

The precise driving force behind rapid browning occurring in the absence of cysteine is unknown and a potential role for cytoplasmic futile creatine cycling warrants further investigation. Previous studies suggested that FGF21 activated by ISR may play a pivotal role in browning^24,67^. Our RNA-seq data reveals that cysteine deficiency induces FGF21 to a greater extent than seen in the tryptophan-free diet +BSO, suggesting that both the ISR and CoA deficiency contribute to FGF21-mediated browning. Mitochondrial stress may hence also induce FGF21, aligning with findings from other reports^68,69^.

The observation that removal of cysteine from the diet does not affect the weight of *Cse* heterozygous mice suggests that the transsulfuration pathway can easily replace dietary cysteine. Targeting cysteine for rapid fat loss may therefore require the inhibition of both the transsulfuration pathway and of cysteine uptake via the cystine/glutamate antiporter (xCT). Developing potent and specific mammalian CSE inhibitors, as previously accomplished for bacterial CSE by our group^70^, would be a significant first step. Given the long-standing interest in developing cysteine uptake inhibitors and cysteine-degrading enzymes in oncology, there are already promising therapeutic candidates such as the xCT inhibitor Erastin and cysteine degrading cysteinase^71,72^. Combining xCT inhibitors with a CSE inhibitor could potentially provide a rapid and innovative solution to combat obesity and other pathologies associated with high cysteine levels, e.g. cystinuria and tumors that require cysteine for growth and progression.

In summary, our findings unravel the profound impact of rendering a non-essential amino acid essential and subsequently removing it from the diet, leading to swift and substantial fat loss. These observations hold significant implications for the field of metabolic medicine, especially in the context of obesity management that could be achieved by manipulating cysteine metabolism.

## Methods

### Mice

Mice were bred and maintained in the Alexandria Center for the Life Sciences animal facility of the New York University School of Medicine, in specific pathogen-free conditions and were fed LabDiet standard 5053 diet prior to experimentation. C57BL/6 mice (Jax 000664), *Gcn2* KO (B6.129S6-Eif2ak4tm1.2Dron/J) and *Fgf21* KO (B6.129Sv(Cg)-Fgf21tm1.1Djm/J) mice were purchased from Jackson Laboratories. *Cse* KO (129/C57BL/6 background) were generated by Rui Wang as previously described and provided by Christopher Hine^13^. Mice in all the experiments were at least 9 weeks old at the starting point of various diets unless described specifically. Het and KOs were co-housed except during food intake measurements and calorie restriction experiments. All animal procedures were performed in accordance with protocols approved by the Institutional Animal Care and Usage Committee of New York University School of Medicine.

### Generation of GDF15 KO mice

GDF15 KO mice (C57BL/6 background) were generated by Genome Editing Shared Resource of Rutgers Cancer Institute. Deletion of GDF15 was performed by coinjecting CRISPR two gRNAs flanking the exon 2 region with Cas9 protein into C57BL/6J zygotes. Verification of CRISPR knockout for the 1.3kb deletion was performed by PCR. Briefly, Primers GDF15A 5’-TCAACTTTAAGCCAGAAGGTGGCG-3’, GDF15B 5’-CTTCGGGGAGACCCTGACTCAGC-3’, and GDF15D 5’-ACTGCGAATCTAGAGAACCCTGAC-3’ were used to amplify the targeted region. Tail snips from founder mice were submitted for Sanger sequencing to confirm homozygous deletion. Founder mouse had a deletion of 1304 bp, which encompasses all of exon 2, the pro-peptide region of GDF15.

### Diets

All custom diets were procured from TestDiet. All diets were based on the defined amino acid diet 5CC7. For individual or dual amino acid depleted diets, the specific amino acid(s) was completely removed, and all other amino acids were increased in proportion. For pantothenic acid-deficient diet, it was removed from the same 5CC7 defined diet. Unless specified control diet refers to the 5CC7 diet. D-Galactose (Sigma G0750) and D-Glucose (Sigma G8270) were purchased from Sigma and were dissolved in water and filter sterilized. A list of diet names is available on request.

### Food intake measurements

Initially, food intake measurements were conducted using metabolic cages; however, mice repeatedly left food debris on the cage bottom compromising the consumption measurements. To address this issue, individually housed mice were provided with a restricted amount of pellets (15-20g), and the remaining food was measured every 24h. Additional intact pellets were added as needed. Mice that left debris on the cage bottom on any day were excluded from the analysis.

### Histology and Immunohistochemistry

Mice were first anesthetized with Ketamine and Xylazine. They were then perfused, first with chilled PBS, followed by 4% PFA in chilled PBS. Tissues were then harvested and fixed for 24 hrs in 4% PFA in PBS at room temperature and then transferred to 70% ethanol. Tissues were then paraffin embedded. Sections (5 µM thick) were cut and stained with H&E. Tissues for immunostaining underwent deparaffinization followed by antigen retrieval for 20 minutes at 100° C with Leica Biosystems ER2 solution (pH9, AR9640) and endogenous peroxidase activity blocking with H_2_O_2_. Sections were incubated with primary antibodies against UCP1 (CST, Cat#: 72298S, Clone: E9Z2V Dilution: 1:1000), CASP3 (CST, Cat#: 9579S, Clone: D3E9 Dilution: 1:200), NQO1 (Sigma-Aldrich, HPA007308, Dilution: 1:100), Custom NRF2 (1:1000, provided by Edward Schmidt, Montana State University)^73^.

Primary antibodies were detected with anti-rabbit HRP-conjugated polymer and 3,3’-diaminobenzidine (DAB) substrate, followed by counter-staining with hematoxylin, all of which were provided in the Leica BOND Polymer Refine Detection System (Cat # DS9800).

### Seahorse Assay

Cells were isolated from inguinal, brachial, and mesenteric lymph nodes. After ACK lysis to remove RBCs, the CD8 T-cell, B-cells and CD11c+ve cells were depleted to leave primarily CD4 T cells. All steps were done at 4° C and used dialyzed fetal bovine serum to avoid amino acids in the serum. Seahorse 24 well plates were plated with Cell-Tak (Corning) as per manufacturer instructions and 100,000 T-cells were plated in 500 µl of room-temperature Seahorse media. After 30 minutes at 37°C, cells were analyzed in Seahorse XFe24 Analyzer.

### Indirect calorimetry

To assess possible changes in metabolic parameters, mice were measured via indirect calorimetry using an open respirometry system (TSE PhenoMaster, TSE Systems GmbH, Germany). Mice were weighed and individually housed in the specialized home cages within a temperature- and humidity-controlled climate chamber (22 ± 0.5 °C, 50 ± 1% relative humidity). The light cycle was set to 12:12 [lights on at 06:30] and the air flow rate for each cage was set to 0.35 l/min (with 0.25 l/min diverted to the gas sensors during the sampling period: 190-s line purge and 10-s active sample). Oxygen consumption [vO_2_ = ml/h], carbon dioxide production [vCO_2_ = ml/h] and, water intake [ml], and activity [beam break counts, X+Y+Z axes] were recorded at 30-min intervals. Respiratory exchange ratio [RER = vCO_2_/vO_2_] were calculated from measured vO2 and vCO2 values. Mice were first acclimated to the metabolic home cages for 24hbefore data was collected for analysis. Mice were housed till set end point of the experiment. Data were acquired and exported with TSE PhenoMaster software V8.1.4.14156 (TSE Systems GmbH, Germany, https://www.tse-systems.com/product-details/phenomaster). After data export, data were uploaded to CalR (https://CalRapp.org/)^74^ for visualization and analysis.

### Preparation of tissue lysates and metabolites analysis

Flash-frozen mouse tissues were homogenized in PBS supplemented with protease (cOmplete, Roche) and phosphatase (PhosSTOP, Roche) inhibitors by IKA Ultra Truemax T8 homogenizer on ice. Aliquots of homogenates were used for RNA isolation immediately. Cells in homogenized tissues were lysed by two freeze thaw cycles and subsequent sonication (Diagenode bioruptor). Protein concentration in the lysates was determined by BCA assay. Aliquots of lysates were filtered through 10K Amicon filters and glutathione and CoA were measured in the flow-through using the following kits: Glutathione detection kit (Cayman 703002) and CoA detection kit (Abcam ab102504). Metabolite concentration was normalized to the protein concentration in lysates. GDF15 and FGF21 were measured in mouse serum by ELISA kits (Abcam ab216947 and R&D Systems MF2100). TG and FAA in mouse serum were analyzed by Cayman (10010303) and Sigma (MAK466) kits respectively.

### Sequencing and differential expression analyses

To study transcriptional response in mouse tissues, total RNA was isolated from the homogenates by trizol (1:10 ratio) method. mRNA was purified by NEBNext® Poly(A) mRNA Magnetic Isolation Module (NEB E7490S) from Turbo-DNAse treated total RNA. A NEB Next® Ultra Library Preparation Kit (NEB E7530S) was used to prepare 0.5 µg of total RNA for RNA-seq. At least three to five animals were used for each experimental condition. The libraries were sequenced using Illumina NextSeq 500 instrument in a paired-end 2×75 cycles setup. The reads were aligned against mouse genome assembly using Hisat2 version 2.1.0. The number of reads in annotated genes was counted using htseq-count version 0.11.0 with option -i set to “gene_id”^75^. The resulting count table was used for differential gene expression analysis with DESeq2 version 1.10.0 using Wald test^76^. Analysis and visualization of the differential expression data was performed with the R software package (version 2.15.1) using the cummeRbund library (version 2.0). Heatmaps were generated based on z-scores of normalized counts utilizing the R library pheatmap. Gene ontology analysis is available at geneontology web applications (http://geneontology.org/).

### Western blotting

Tissue lysates supplemented with LDS loading buffer and 10 mM DTT were heated up for 5 minutes at 95°C. Proteins were separated on Bis-Tris SDS-PAGE, transferred onto nitrocellulose membrane and probed with anti-Phospho-eIF2α (Cell Signaling Technology 3597), anti-eIF2α (Cell Signaling Technology 2103) anti-Phospho-ACC (Cell Signaling Technology 3661), anti-ACC (Cell Signaling Technology 3662), anti-β-Tubulin (Proteintech 10068-1-AP) and anti-β-Actin−Peroxidase antibody (Sigma A3854).

### Mass-spectrometry

#### Sample Preparation Stool Samples

Stool samples and food pellets were weighed into bead blaster tubes containing zircon beads. Extraction buffer containing 80% methanol with 500nM Metabolomics amino acid standard mix (Cambridge Isotopes Laboratory, MA) was added to each to reach a final concentration of 10mg/mL. Samples were homogenized using D2400 BeadBlaster homogenizer (Benchmark Scientific, NJ) then spun at 21kg for 3min. Then, 450uL of metabolite extract was transferred to a new 1.5mL Eppendorf tube and dried using Speedvac. Samples were reconstituted in 50uL of MS grade water and sonicated for 2 mins. Then, samples were spun at 21,000 g for 3 mins. Samples were transferred to glass LC vials for analysis by LCMS.

#### Liver Samples

Approximately 300 mg of liver was homogenized in 1 mL of PBS and then subjected to three freeze-thaw cycles. Liver samples were filtered using a 10kDa filter. Protein concentrations prior to deprotenization were measured using BSA standard curve and found to be between 12.8 – 42.0 mg/mL. For metabolomics extracts, on average protein concentration was determined to be 28.58 mg/mL per 300 mg of tissue. This value was used to scale all liver extracts to each other. Scaled liver extracts were transferred to bead blaster tubes with zircon beads and homogenized using D2400 BeadBlaster homogenizer (Benchmark Scientific, NJ) in cold 80% methanol spiked with 500 nM Metabolomics amino acid standard mix (Cambridge Isotopes Laboratory, MA). Samples were centrifuged at 21,000 g for 3 min to pellet any insoluble materials. Then, 450 uL of metabolite extract was transferred to a new 1.5mL Eppendorf tube and dried down using Speedvac. Samples were reconstituted in 50uL of MS grade water and sonicated for 2 mins. Then, samples were spun at 21,000 g for 3 min. Samples were transferred to glass LC vials for analysis by LCMS.

#### Urine Samples

Urine samples were collected and stored as frozen aliquots between ∼5-7µL. For metabolite extraction, 5µL of urine was transferred to a glass insert and extracted using 195µL of cold 80% methanol spiked with 500nM Metabolomics amino acid standard mix (Cambridge Isotopes Laboratory, MA). Glass inserts were transferred into 1.5mL Eppendorf tubes and spun at 3 kg for 10 min to pellet insoluble material. Then, 180 µL of extract was transferred to a 1.5mL Eppendorf tube and dried down completely using Speedvac. To each, 20 µL of MS grade water was added to reconstitute metabolites. Samples were sonicated for 2 mins and then centrifuged for 3 min at 21kg. Then 15 µL of samples was transferred to a glass LCMS vial for analysis.

#### LC-MS/MS analysis

LC-MS/MS with the hybrid metabolomics method – Samples were subjected to an LCMS analysis to detect and quantify known peaks. A metabolite extraction was carried out on each sample based on a previously described method1. The LC column was a MilliporeTM ZIC-pHILIC (2.1 x150 mm, 5 μm) coupled to a Dionex Ultimate 3000TM system and the column oven temperature was set to 25°C for the gradient elution. A flow rate of 100 μL/min was used with the following buffers; A) 10 mM ammonium carbonate in water, pH 9.0, and B) neat acetonitrile. The gradient profile was as follows; 80-20%B (0-30 min), 20-80%B (30-31 min), 80-80%B (31-42 min). Injection volume was set to 2 μL for all analyses (42 min total run time per injection).

MS analyses were carried out by coupling the LC system to a Thermo Q Exactive HFTM mass spectrometer operating in heated electrospray ionization mode (HESI). Method duration was 30 min with a polarity switching data-dependent Top 5 method for both positive and negative modes. Spray voltage for both positive and negative modes was 3.5kV and capillary temperature was set to 320°C with a sheath gas rate of 35, aux gas of 10, and max spray current of 100 μA. The full MS scan for both polarities utilized 120,000 resolution with an AGC target of 3e6 and a maximum IT of 100 ms, and the scan range was from 67-1000 m/z. Tandem MS spectra for both positive and negative mode used a resolution of 15,000, AGC target of 1e5, maximum IT of 50 ms, isolation window of 0.4 m/z, isolation offset of 0.1 m/z, fixed first mass of 50 m/z, and 3-way multiplexed normalized collision energies (nCE) of 10, 35, 80. The minimum AGC target was 1e4 with an intensity threshold of 2e5. All data were acquired in profile mode.

#### Hybrid Metabolomics Data Processing

Relative quantification of metabolites – The resulting ThermoTM RAW files were converted to mzXML format using ReAdW.exe version 4.3.1 to enable peak detection and quantification. The centroided data were searched using an in-house python script Mighty_skeleton version 0.0.2 and peak heights were extracted from the mzXML files based on a previously established library of metabolite retention times and accurate masses adapted from the Whitehead Institute^77^ and verified with authentic standards and/or high resolution MS/MS spectral manually curated against the NIST14MS/MS^78^ and METLIN (2017)^79^ tandem mass spectral libraries. Metabolite peaks were extracted based on the theoretical m/z of the expected ion type e.g., [M+H]+, with a ±5 part-per-million (ppm) tolerance, and a ± 7.5 second peak apex retention time tolerance within an initial retention time search window of ± 0.5 min across the study samples. The resulting data matrix of metabolite intensities for all samples and blank controls was processed with an in-house statistical pipeline Metabolize version 1.0 and final peak detection was calculated based on a signal to noise ratio (S/N) of 3X compared to blank controls, with a floor of 10,000 (arbitrary units). For samples where the peak intensity was lower than the blank threshold, metabolites were annotated as not detected, and the threshold value was imputed for any statistical comparisons to enable an estimate of the fold change as applicable. The resulting blank corrected data matrix was then used for all group-wise comparisons, and t-tests were performed with the Python SciPy (1.1.0)^80^ library to test for differences and to generate statistics for downstream analyses. Any metabolite with p-value < 0.05 was considered significantly regulated (up or down). Heatmaps were generated with hierarchical clustering performed on the imputed matrix values utilizing the R library pheatmap (1.0.12, available at https://CRAN.R-project.org/package=pheatmap). Volcano plots were generated utilizing the R library, Manhattanly (0.2.0). In order to adjust for significant covariate effects (as applicable) in the experimental design the R package, DESeq2 (1.24.0)^76^ was used to test for significant differences. Data processing for this correction required the blank corrected matrix to be imputed with zeroes for non-detected values instead of the blank threshold to avoid false positives. This corrected matrix was then analyzed utilizing DESeq2 to calculate the adjusted p-value in the covariate model.

### AAV-mediated rescue experiments

Mice were injected retro-orbitally with 2*10^11^ liver specific AAV8 TBG-CSE or AAV8 TBG-eGFP viral particles per mouse (Vector Biolabs). Two weeks after AAV administration, mice we placed on a No Cys diet and after one-week tissues and serum were harvested and analyzed.

### Heavy glucose tracing

Mice were fasted for 18 hours after 3 days of No Cys diet. They were then orally given 1.25 mg/kg of D-Glucose-13C6 (Sigma) dissolved in water. Mice were euthanized at 45 minutes to collect livers or at 2 hours to collect urine. Liver was processed as described in the “Preparation of tissue lysates and metabolites”. Subsequently liver lysates and urine were analyzed as described in the “Mass spectrometry” section.

The data were then processed and naturally expected levels of each molecule’s isotope were subtracted by the expected natural frequency of each isotope multiplied by each individual sample’s lowest weight isotope amount for the molecule.

These corrected data were then used for the analysis and is included in Supplementary Table 6.

## Author Contributions

A.V., I.G., E.N. and D.R.L. conceived the project, A.V. and I.G. designed experiments and analyzed data. A.V., I.G., B.G.L., D.D., Y.M., I.S., M.P., R.J., D.J., A.C.M., performed experiments. M.G-J., E.P.W., R.W. and T.P. provided resources, T.P. and M.E.P. advised experiments, A.V., I.G., D.R.L and E.N. wrote the manuscript with input from all the authors. D.R.L and E.N. acquired funding and supervised the research.

## Acknowledgements

We thank the members of the Nudler and Littman laboratories for their valuable discussions and critical review of the manuscript. We thank Klaudia Laborc of the Mar lab and NYU Rodent Behavior Laboratory (RBL) for assistance with the metabolic cage experiments, Edward Schmidt (Montana State University) for the custom NRF2 antibody, S. Y. Kim and the NYU Rodent Genetic Engineering Laboratory (RGEL) for rederivation of mutant mice and C. Loomis and the Experimental Pathology Research Laboratory of NYULMC for histology and Immunohistochemistry. We also thank Susan Gottesman for editing the manuscript. The Experimental Pathology Research Laboratory is supported by National Institutes of Health Shared Instrumentation grants S10OD010584-01A1 and S10OD018338-01. This work was supported by the NSERC RGPIN-2023-05099 (R.W.), Ludwig Princeton Branch, and Cancer Grand Challenges (NCI 1OT2CA278609-01, CRUK (CGCATF-2021/100022) to E.W., Blavatnik Family Foundation (E.N.); Howard Hughes Medical Institute (E.N., D.R.L); and NIH R01AI158687 (D.R.L.).

## Conflicts of Interest

D.R.L consults for and has equity interest in Vedanta Bioscience, Sonoma Immunotherapeutics, Immunai, IMIDomics, and Pfizer, Inc.

---

**Extended Data Fig. 1:**
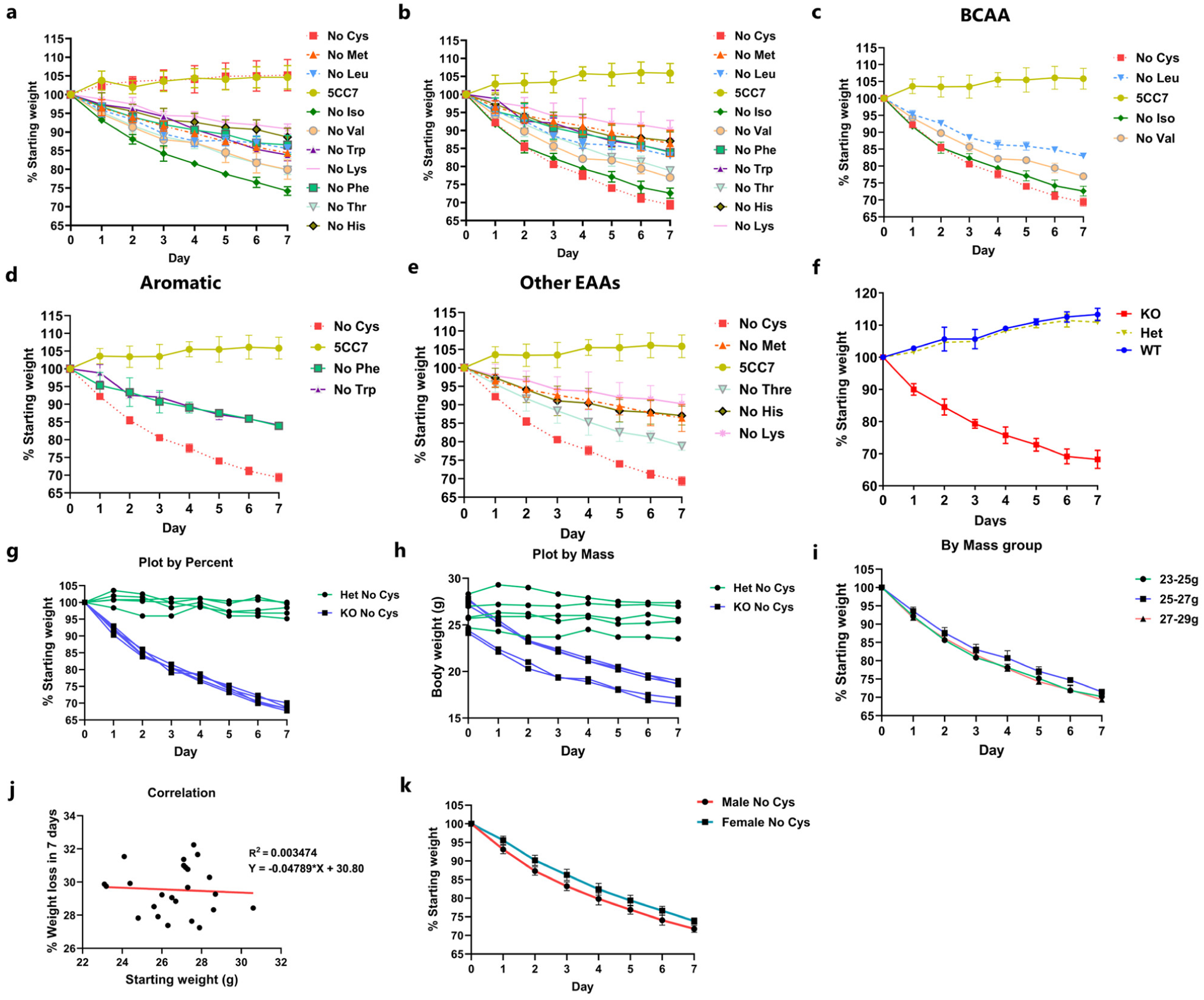
Further characterization of cysteine-induced weight loss. **a,** Het and **b,** KO weight curves with various essential amino acid deficiencies (n>=4). **c-e**, KO on various essential amino acid deficiencies by class: **c**, Branched chain amino acids; **d**, Aromatic amino acids; **e**, Other EAAs. **f**, Weight curves of WT compared to *Cse* Het and KO mice on No Cys diet. Weight loss curves of *Cse* KO on No Cys plotted individually by **g,** percentage **h,** mass **i**, Weight loss curves by percentage plotted according to different starting weights (n≥5 in all groups) **j**, Percentage weight loss at Day 7 on a No Cys diet for *Cse* KO plotted against starting mass **k**, Weight loss curves of male vs female *Cse* KO mice on a No Cys diet (n=9)

**Extended Data Figure 2:**
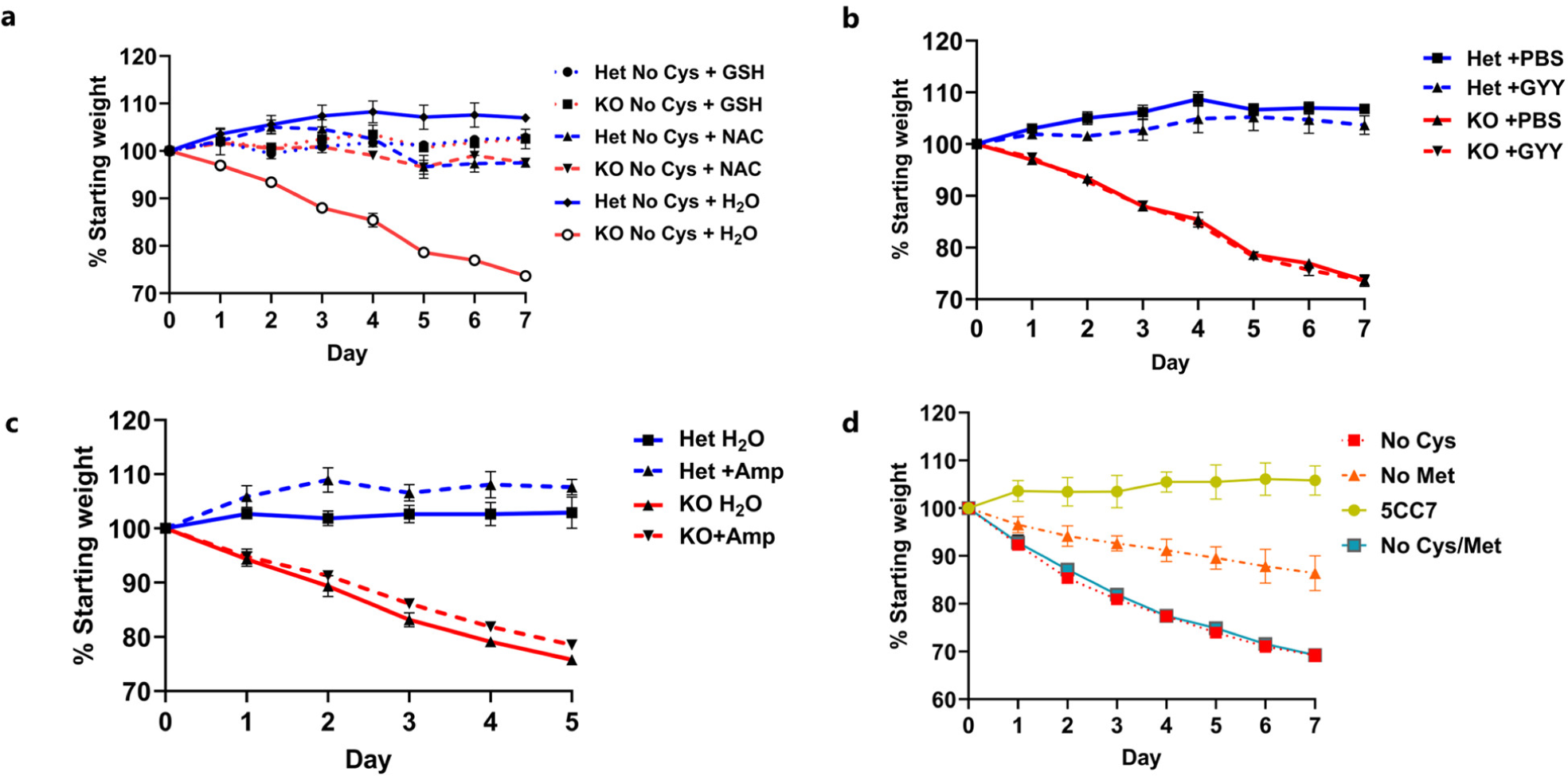
Effect of other sulfur containing molecules on weight loss. **a**, Weight curves of Het and KO mice on a No Cys diet with either 0.4% NAC or 1% GSH in drinking water (n=4). **b**, Weight curves of mice on a No Cys diet with or without daily injections of 40 mg/kg of GYY4137 (n=2). **c**, Weight curves on a No Cys diet with and without antibiotic treatment (1 g/L Ampicillin) (n=3). **d**, Weight curves of mice on a No Met No Cys (Het) compared to No Cys, No Met, or control diets (*Cse* KO) (n=4). Hets and KOs mice were co-housed for each experimental condition.

**Extended Data Figure 3:**
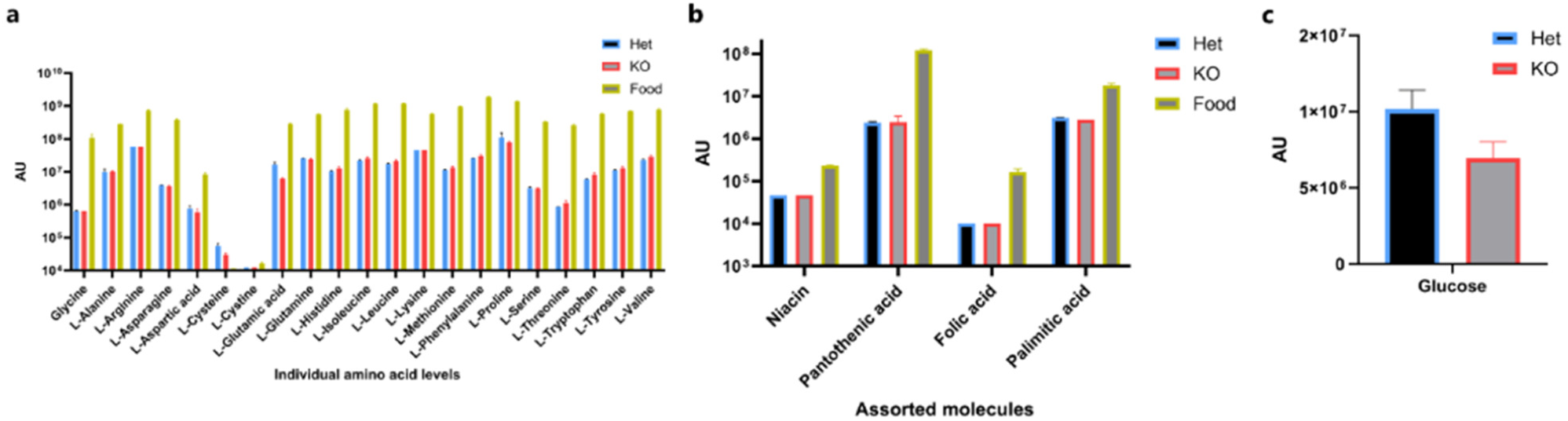
Nutrient absorption is not affected by No Cys diet in *Cse* KO. **a**, Amino acid levels in the administered diet and in Het or KO stool from microbiota-depleted mice fed No Cys diet for 3 days. **b**, Vitamins and palmitic acid levels **c**, Glucose levels in stools of mice on Day 3 of No Cys diet (n=3).

**Extended Data Fig. 4:**
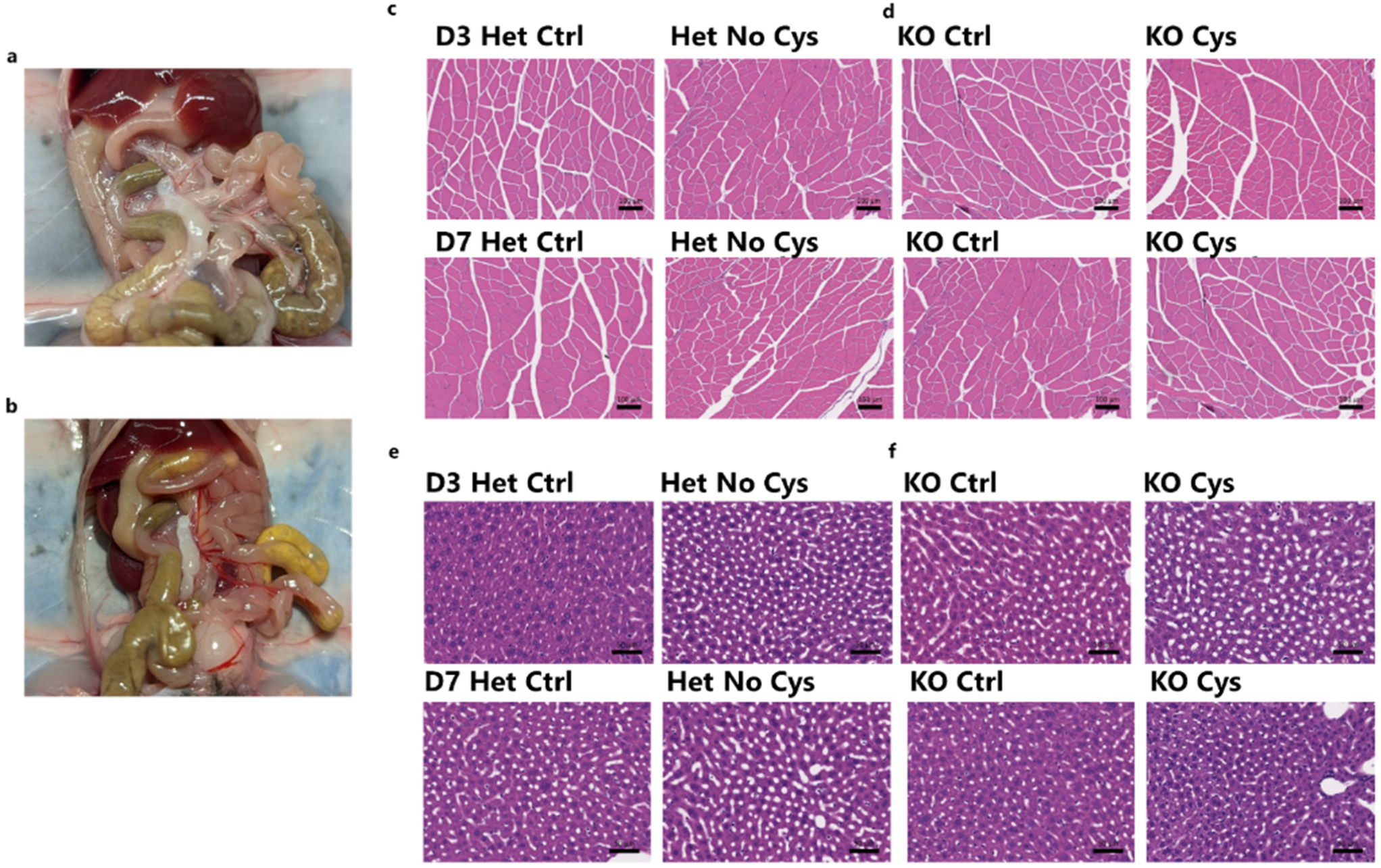
Gross histology of skeletal muscle and liver. **a,b**, Representative visceral fat image of Het (a) and KO (b) mice on No Cys diet after 3 days (n=4). **c-f,** Representative H&E staining of skeletal muscle (quadriceps) (c,d) and liver (e,f) from Het (c,e) and KO (d,f) mice on Days 3 and 7 of calorically restricted Ctrl or No Cys diets (n=4).

**Extended Data Fig. 5:**
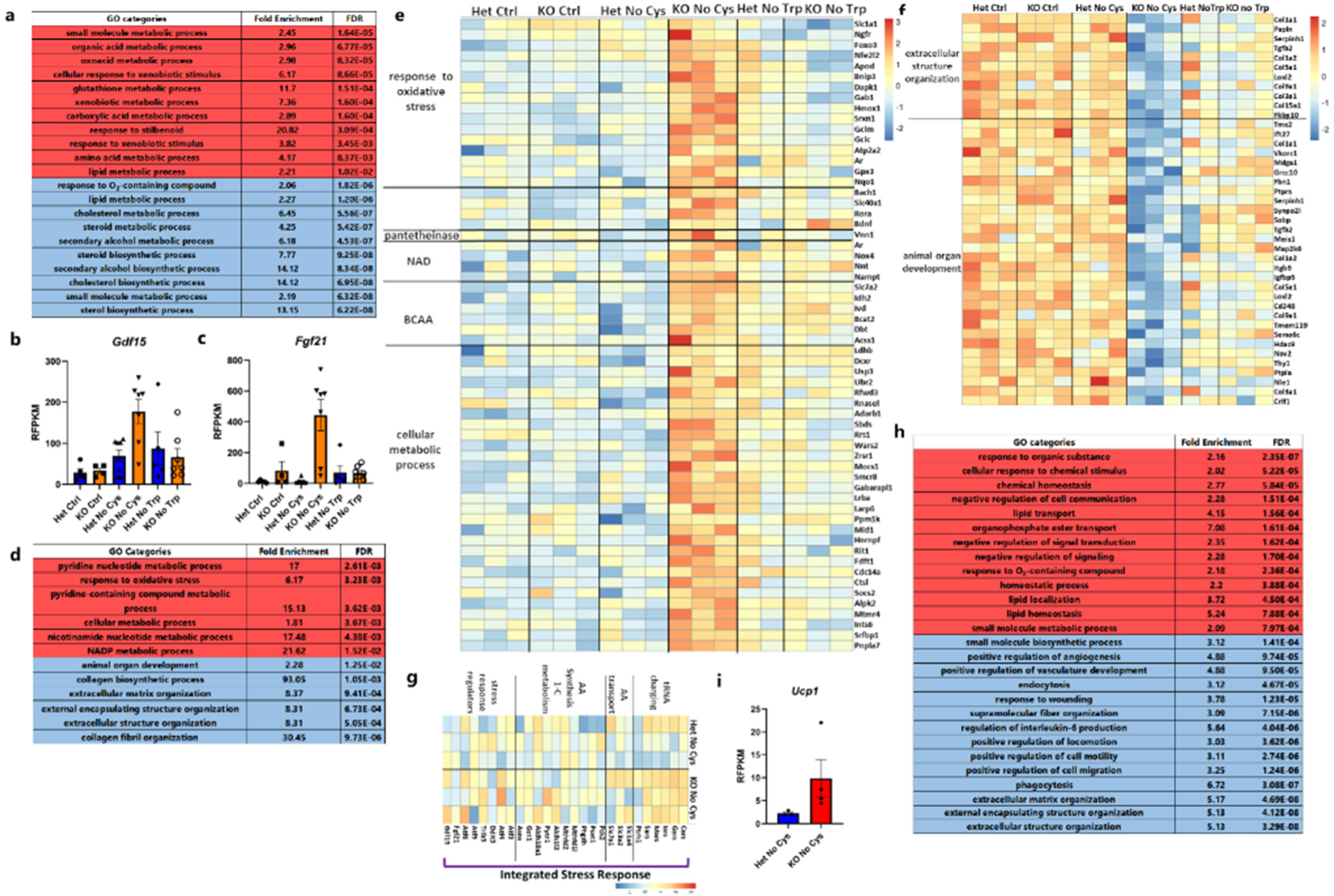
Transcriptional response to cysteine deficiency. **a**, GO enrichment categories from liver comparing Het and KO mice on No Cys diet. **b**, *Gdf15* RFPKM values from liver across all groups. **c**, *Fgf21* RFPKM values from liver across all groups. **d**, GO enrichment categories from muscle comparing Het and KO mice on No Cys diet. **e-g**, Muscle bulk RNA-Seq data represented as heatmap for genes in oxidative stress, cellular metabolic process, branched chain amino acid metabolism, NAD, and pantetheinase that are upregulated (e), for genes in extracellular organization and animal organ development that are specifically downregulated (f), and for genes in the ISR that appear minimally affected (g) in No Cys KO mice at Day 2. **h**, GO enrichment categories from epididymal fat pad comparing Het and KO mice on No Cys diet. **i**, *Ucp1* RFPKM values from epididymal fat pad. All data are at Day 2 based on mice shown in the RNA-seq schematic in Figure 3a. For GO enrichment table, Red indicates upregulated in KO and blue indicates downregulated.

**Extended Data Fig. 6:**
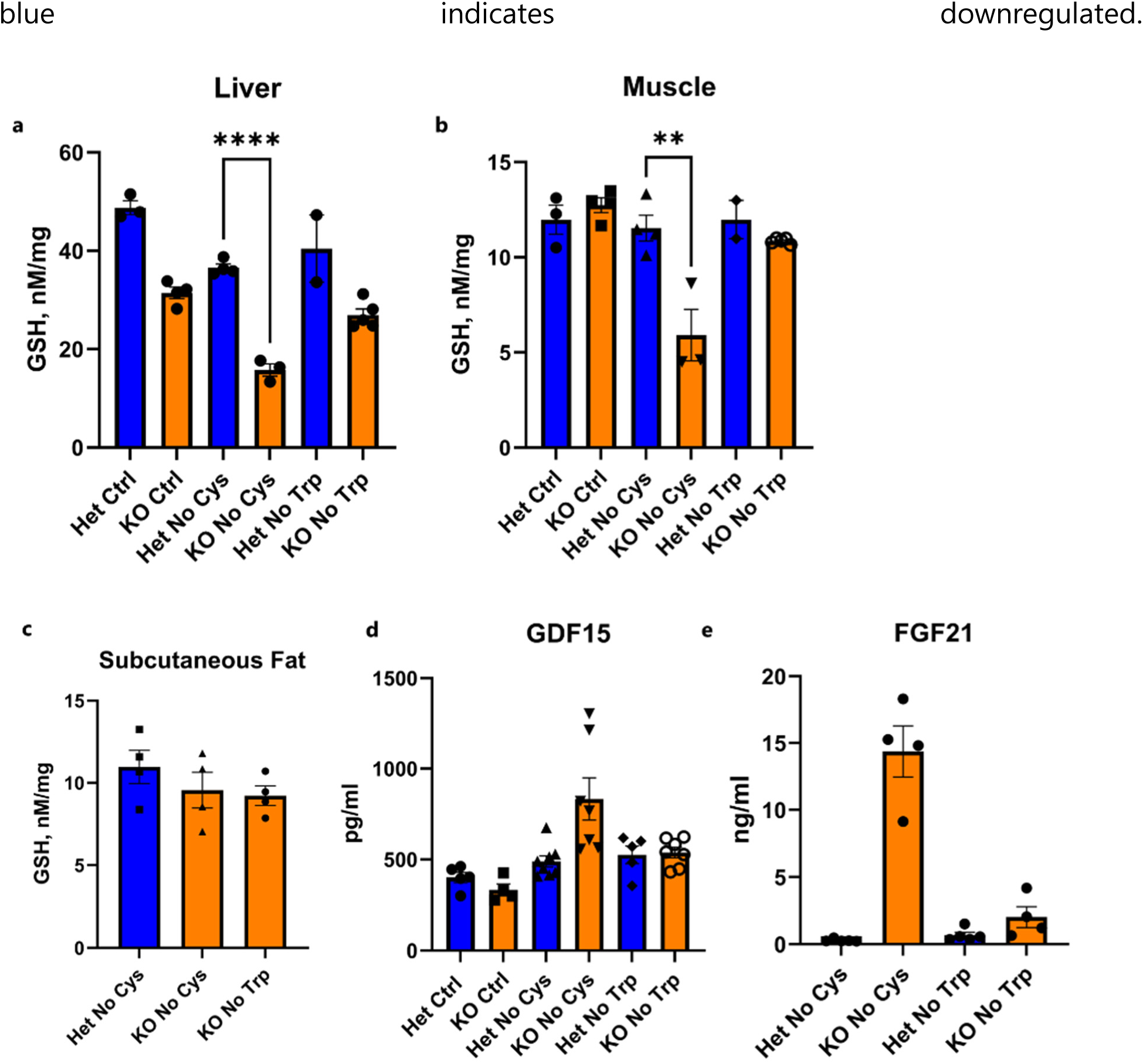
Effect of cysteine restriction on tissue GSH, serum GDF15 and FGF21 levels. **a-e,** GSH levels in liver (a), muscle (b) and subcutaneous fat pad (c) GDF15 (d) and FGF21 (e) levels in serum from *Cse* Het and KO mice on Day 2 of CR diets without Cys or Trp or Ctrl diet.

**Extended Data Figure 7:**
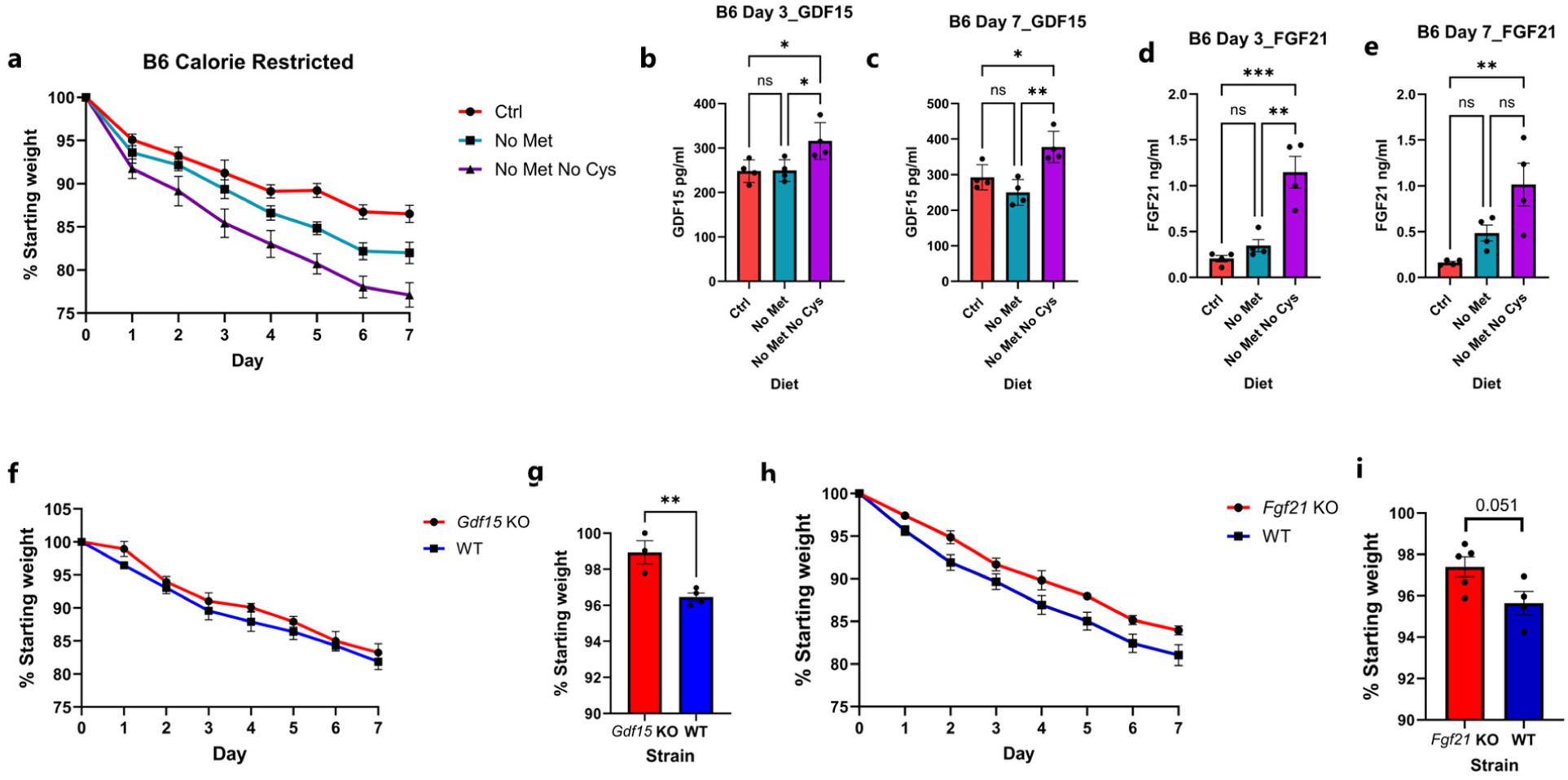
Effect of Cysteine and methionine dual restriction on wild type C57BL/6 mice. **a**, Weight loss curves of calorie restricted (2.1 g/day) male mice on Ctrl, No Met or No Met No Cys diet (n=4). **b,** Serum GDF15 levels across all three conditions on Day 3 or **c**, Day 7. Serum FGF21 levels across all three conditions on **d**, Day 3 or **e**, Day7 (n=4). **f**, Weight loss curves of male *Gdf15* KO or WT mice on No Met No Cys diet (n≥3) **g**, Weight after one day of No Met No Cys diet in *Gdf15* KO or WT mice **h**, Weight loss curves of female *Fgf21* KO or WT mice on No Met No Cys diet (n≥4) **i**, Weight after one day of No Met No Cys diet in *Fgf21* KO or WT mice.

**Extended Data Figure 8:**
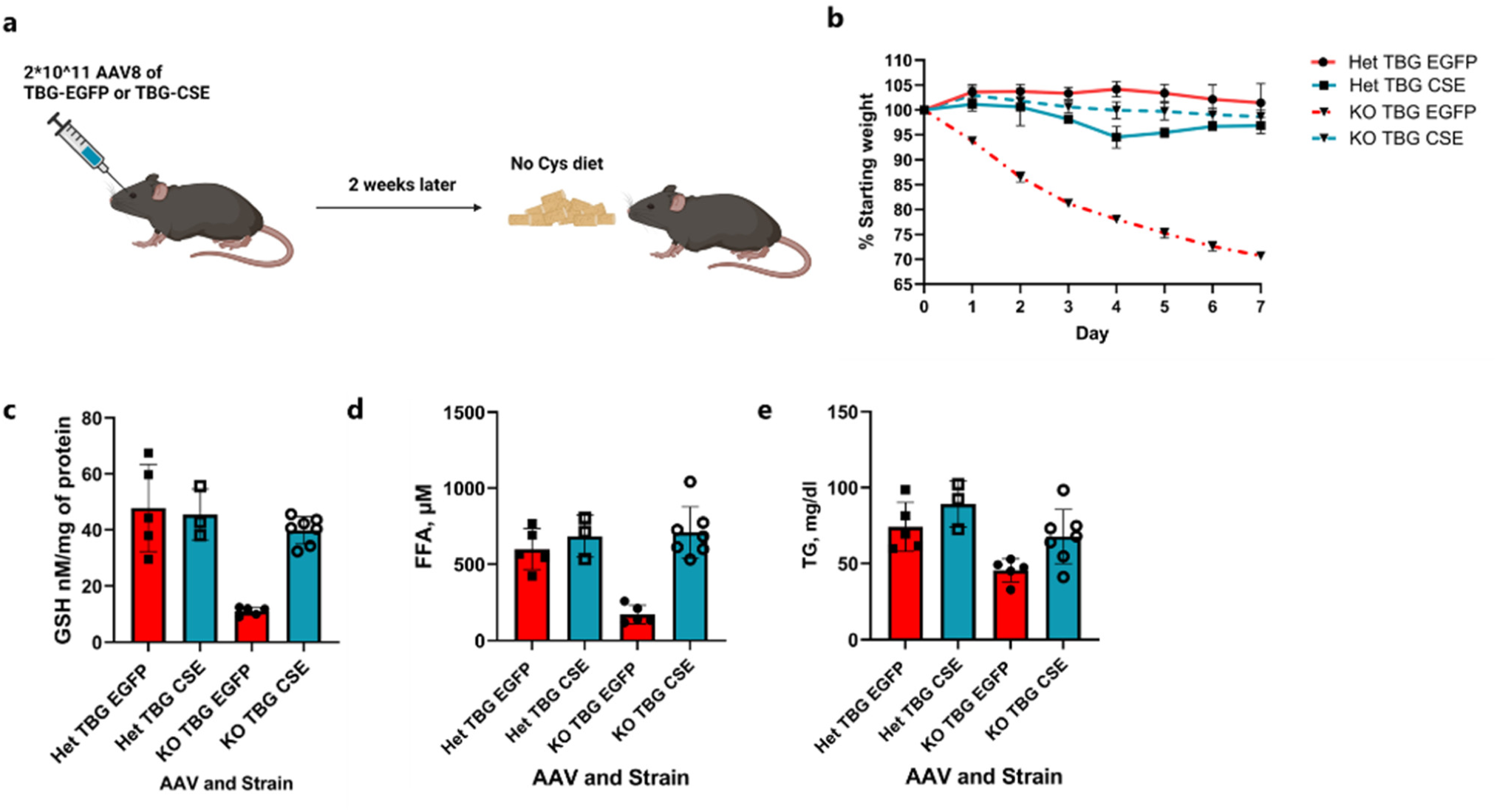
Effect of liver specific *Cse* expression on rescuing weight loss in cysteine-deficient mice. **a**, Experimental scheme. **b,** Weight loss curves of *Cse* Het or KO mice infected with AAV8-TBG-EGFP or AAV8-TBG-CSE on a No Cys diet. **c-e**, Levels at day 7 of GSH (d) and FFA (e) in liver and TG in serum (f) across all 4 groups (n≥3).

**Extended Data Fig. 9:**
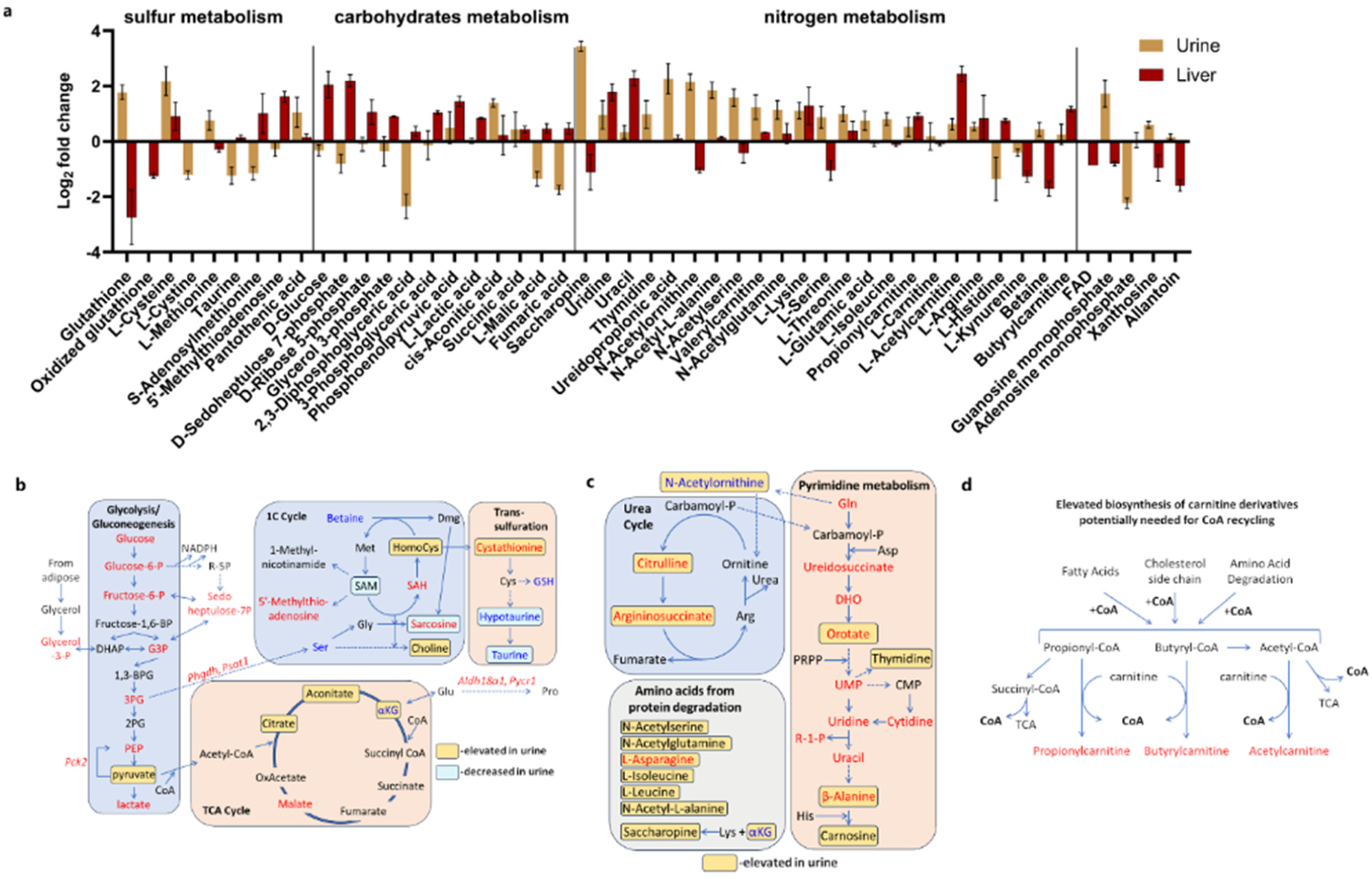
Metabolite differences in liver and urine of *Cse* Het and KO mice on No Cys diet. **a**, Select metabolites from liver (n=3) and urine (n>=6) of KO mice, normalized to levels in Het mice. **b**, Glycolysis and TCA cycle pathway metabolites represented with data from liver (n=3) and urine (n>=6) **c**, Urea cycle and pyrimidine metabolism pathway metabolite profiles represented with data from liver (n=3) and urine (n>=6). Red is higher in KO and Blue is lower. Yellow and blue boxes indicate metabolites that are elevated or reduced, respectively, in urine of KO mice. **d**, Carnitine and CoA metabolism model to explain excess Acyl-carnitines detected in urine.

**Extended Data Figure 10:**
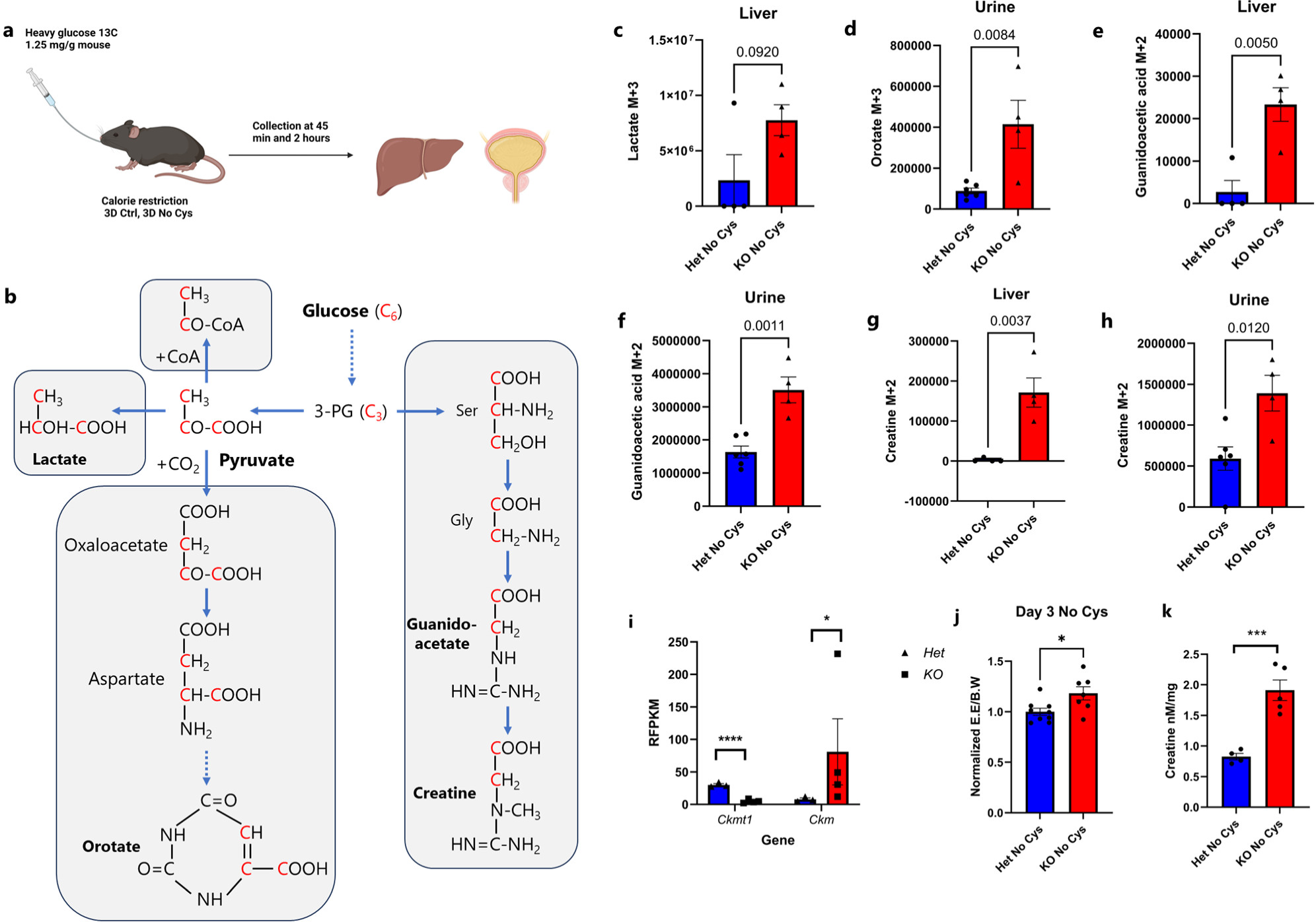
^13^C-Glucose tracing in *Cse* het and KO mice on Cys-deficient CR diet. **a,** Scheme of ^13^C glucose tracing experiment (n=4 for liver, n≥4 for urine, all males). **b,** Pathway depicting the formation of heavy carbon labeled metabolites from m+6 glucose. **c,** Levels of m+3 lactate in *Cse* Het or KO mice at 45 minutes in liver **d,** Levels of m+3 orotate in Urine at 2h in *Cse* Het or KO mice. **e** and **f**, Levels of guanidoacetate m+2 in liver (e) at 45 minutes and urine (f) at 2h in Het or KO mice. **g** and **h**, Levels of creatine m+2 in liver (g) at 45 minutes and urine (h) at 2h in Het or KO mice. **i,** Expression of *Ckmt1* and *Ckm* in epididymal fat of *Cse* Het and KO mice on No Cys diet (see also Supplementary table 4) (n≥3) **j,** Peak energy expenditure (EE) at Day 3 of a No Cys diet normalized to body weight, subsequently normalized to Het data (n≥7). **k,** Creatine levels in liver at Day 6 on a No Cys diet in Het and KO mice (n≥4).

**Extended Data Fig. 11:**
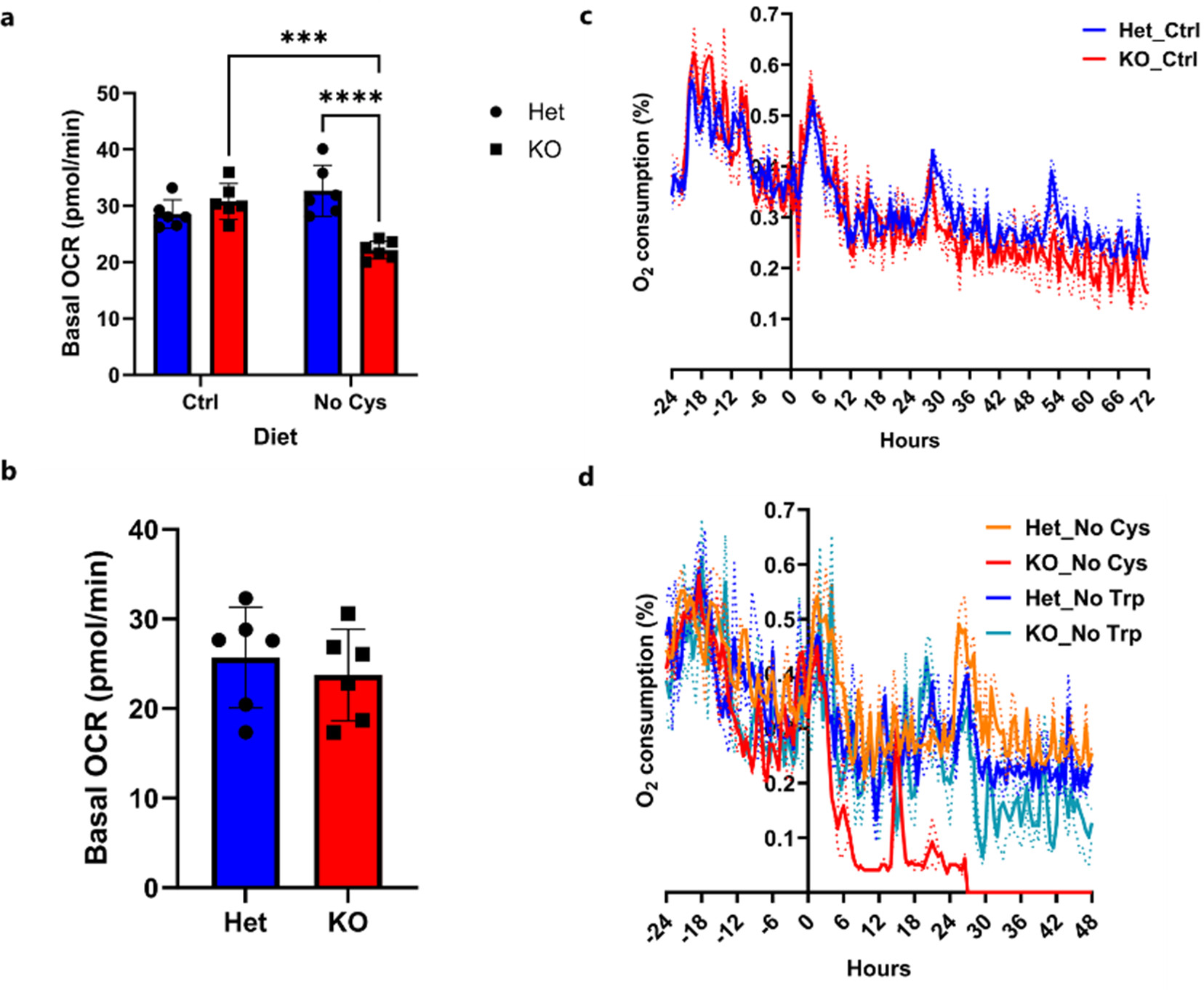
*Cse* KO mice become more dependent on glycolysis when subjected to a No Cys diet. Seahorse cell energy phenotype assay and basal OCR of CD4 T cells isolated from *Cse* Het and KO mice on **a**, Ctrl or No Cys diet **b**, No Trp diet for 7 days. **c, d,** O_2_ consumption of *Cse* Het and KO mice restricted to 10% galactose as energy source and fed a (c) control diet (n=4) or (d) a No Cys diet compared to a No Trp diet (n=2) as per scheme in Fig 5g.

**Extended Data Fig. 12:**
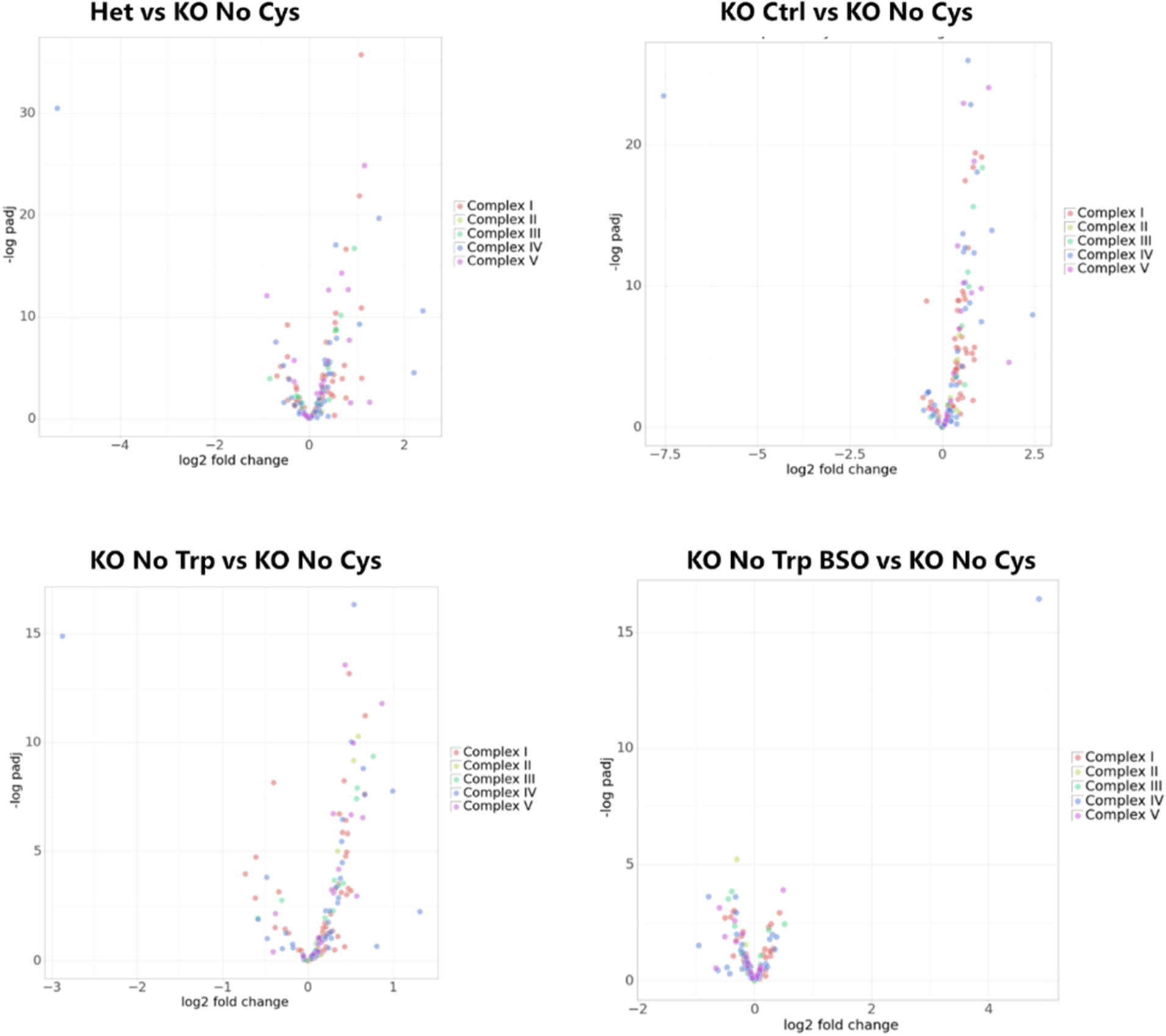
Volcano plots of mitochondrial complex genes across both RNA-seq experiments. Data points are based on liver bulk RNA-Seq data shown in Figures 3 and 4 (n>=3). Colors represent genes encoding proteins in each of the complexes of the electron transport chain.

**Extended Data Figure 13:**
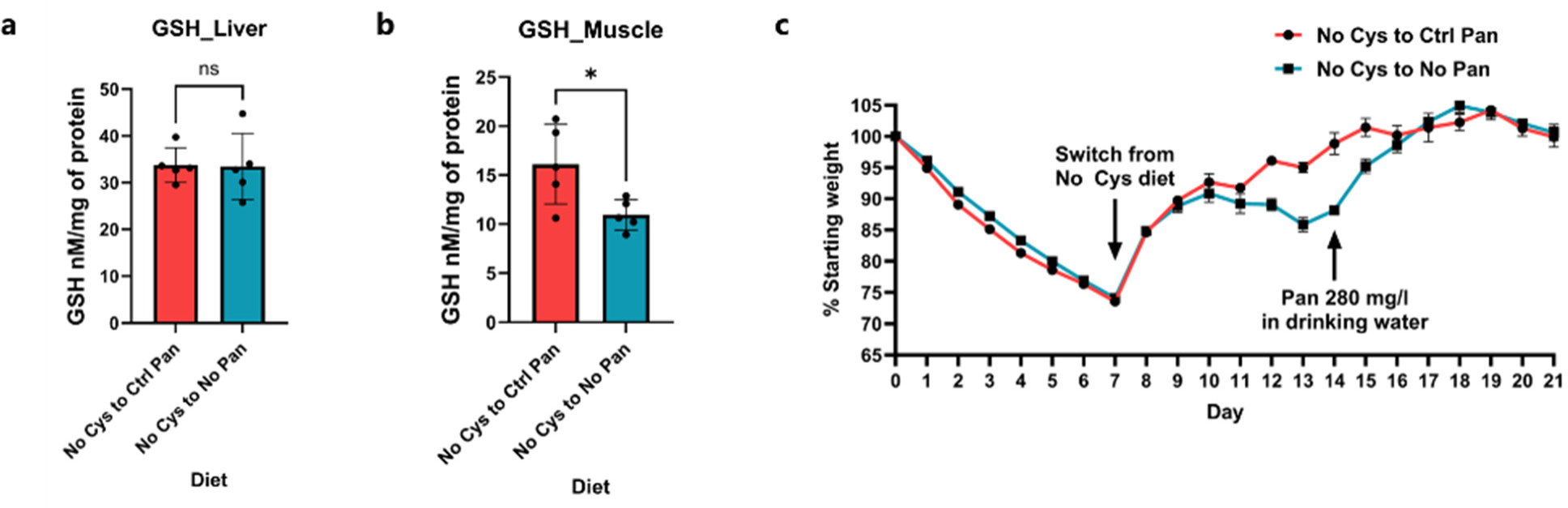
Role of CoA in weight recovery. **a**, Liver and **b,** Muscle GSH levels at Day 14 of experiment in Fig 5l (n=5) **c**, Weight loss curves of female *Cse* KO mice on No Cys diet for 7 days, followed by either 7 days on Ctrl diet or Vit-B5 (Pan) free diet followed by 7 days on a Ctrl diet or a Vit-B5 free diet with supplemented with 280 mg/ml of Vit-B5 (Pan) (n=4).

